# Using temperature to analyse the neural basis of a time-based decision

**DOI:** 10.1101/2020.08.24.251827

**Authors:** Tiago Monteiro, Filipe S. Rodrigues, Margarida Pexirra, Bruno F. Cruz, Ana I. Gonçalves, Pavel E. Rueda-Orozco, Joseph J. Paton

**Affiliations:** Champalimaud Centre for the Unknown, Lisbon, Portugal; Department of Biology, University of Oxford, UK; Sainsbury Wellcome Centre for Neural Circuits and Behaviour, University College London, London, UK; NeuroGEARS Ltd., London, UK; Institute of Neurobiology, UNAM Juriquilla, Mexico

## Abstract

The basal ganglia (BG) are thought to contribute to decision-making and motor control by influencing action selection based on consequences. These functions are critically dependent on timing information that can be extracted from the evolving state of neural populations in the striatum, the major input area of the BG. However, it is debated whether striatal activity underlies latent, dynamic decision processes or kinematics of overt movement. Here, we measured the impact of temperature on striatal population activity and the behavior of rats and compared the observed effects to neural activity and behavior collected in multiple versions of a temporal categorization task. Cooler temperatures caused dilation, and warmer temperatures contraction, of both neural activity and patterns of judgment in time, mimicking endogenous decision-related variability in striatal activity. However, temperature did not similarly affect movement kinematics. These data provide compelling evidence that the time course of evolving striatal population activity dictates the speed of a latent process that is used to guide choices, but not moment by moment kinematics. More broadly, they establish temporal scaling of population activity as a likely cause and not simply a correlate of timing behavior in the brain.

## INTRODUCTION

Much of behavior is dependent on time. Humans and other animals must extract temporal structure from the environment to learn to anticipate events, to understand relationships between actions and consequences, and estimate time, implicitly or explicitly, to plan and properly sequence and coordinate action. For tasks as varied as waiting at a stoplight to a hummingbird foraging for nectar, time is fundamental.

Timing mechanisms appear to be distributed across the nervous system, reflecting the importance of time information for much of brain function^1,2^. However, one common requirement among diverse functions is the need to create an index, ordering and spacing information along the temporal dimension such that useful relations can be extracted and outputs appropriately coordinated. On the scale of seconds to minutes at which much of behavior unfolds, neuronal population dynamics represent a candidate means of both encoding and generating temporal patterns. Artificial neural network models have explored evolving population activity as a basis for timing sensory events^3^ and movements^4^. And correlations between behavior and the time course of population activity have lent some support to the hypothesis that time-varying patterns of activity within a population perform temporal computations^5–8^. However, correlations do not imply causation. The critical prediction of these “population clock” hypotheses is that slowing or speeding the temporal evolution of activity should cause a corresponding dilation or contraction of the temporal functions performed using that activity.

One brain system where time information appears to be critical is the basal ganglia (BG), an evolutionarily ancient set of brain structures thought to contribute to appropriate action selection based on experience. A dominant view holds that the BG embed core features of reinforcement learning (RL)^9^ algorithms. In mammals, inputs from a diverse set of territories in cortex, thalamus, and limbic brain structures convey information about the state of the world that converges with dense dopaminergic innervation in the major input area of the BG, the striatum. The input from dopamine neurons is thought to teach striatal circuits about the value of taking particular actions in a given state, information that can ultimately be conveyed to downstream brainstem and thalamo-cortical motor circuits to bias selection or otherwise specify features of actions. To accomplish such a function, the BG would need access to information about ordering and spacing along the temporal dimension, either implicitly or explicitly, both to extract meaningful relations between the environment, actions and outcomes that drive learning^10,11^, and to coordinate the production of actions in time^12,13^. Interestingly, data from people with BG disorders^14,15^ and human fMRI^16,17^ have consistently identified the BG as being involved in timing behavior. In addition, lesions and pharmacological manipulations of the striatum can cause deficits in temporal estimation and reproduction^18,19^. Lastly, recordings from striatal populations have demonstrated that time information can be decoded from neural activity, and this information correlates with variability in timing behavior^7,20–23^. Specifically, the state of striatal population activity continuously changes along reproducible trajectories during behavioral tasks that require time estimation, advancing more quickly when animals report long judgments, and more slowly when they report short judgments^4,7,19,23^.

To test whether variability in the speed of BG population dynamics merely correlates with or directly regulates timing function, we sought to experimentally manipulate dynamics as animals reported temporal judgments. Interestingly, despite being composed of elements with differing temperature dependencies (e.g., ion channel conductances and synaptic transmission)^24^, neural circuits in at least some systems can produce patterns of activity that systematically slow down or speed up with decreasing or increasing temperature^25,26^. For this reason, temperature manipulations offer a potential method to test hypotheses regarding the relationship between the speed of neural dynamics and function^27^. Indeed temperature manipulations in the zebra finch have been used to identify area HVC as a locus within the song production circuit that contributes to the temporal patterning of bird song^28^. Similar temperature manipulations in humans have identified a subregion of speech motor cortex that regulates the speed of speech^29^, a region of rodent medial frontal cortex that controls the timing of a delayed movement^30^, and cooling the medial septum has been shown to slow theta oscillations in the hippocampus^31^. However, temperature can have distinct effects on neural activity depending on cellular and circuit-level characteristics of the area in question^28,32^.

Here, we used a custom thermoelectric device (TED) to systematically vary the temperature of striatal tissue, both warming and cooling relative to a baseline condition. We found that temperature affected overall activity levels non-monotonically, with both warming and cooling relative to a control temperature producing lower baseline firing rates overall. In contrast, temperature manipulations caused monotonic and graded changes in the temporal scaling of neural population activity, mimicking decision-related variability in activity observed during multiple versions of a temporal judgment task. Temperature manipulations also caused bidirectional and graded changes in animals’ timing judgments, mirroring the temperature-dependent modification of temporal scaling of neural activity as well as the observed relationship between temporal scaling of activity and animals’ judgments. Strikingly, these results were not accompanied by similar effects of temperature on movement execution. Instead, though more modest than the effects of temperature on animals’ judgments, the pattern of average speed of animals’ movements was a non-monotonic function of temperature, similar to the observed non-monotonic effect of temperature on baseline firing rate. Together, these findings imply that distinct aspects of behavior may be controlled by temporal evolution of population activity and the overall levels of activity. Such distinctions suggest that continuously evolving patterns of activity in the striatum support the placement of discrete behavioral transitions in time, as required for action selection and decision-making, but do not provide moment by moment motor command-related signals required for continuous control. Instead of the BG providing high dimensional continuous control signals, overall activity appears to provide a low dimensional gain signal applied to motor programs executed elsewhere, consistent with the role of BG circuits in modulating movement vigor.

## RESULTS

### Striatal temperature modified neural population speed monotonically, and baseline activity levels non-monotonically

Temperature has been shown in some systems to alter the speed of neural population activity while maintaining its general pattern^25,26^. Thus, we first examined the effect of temperature on striatal neural activity.

First, we developed a thermoelectric device (TED)^33^ (**Fig. 1A**) based on the Peltier effect and used it to achieve closed-loop control over the temperature of silver metal probes implanted in brain tissue. To characterize the spatio-temporal profile of temperature changes in the brain, we measured temperature at different distances from the tip of a probe implanted in dorsal striatum (DS), setting our TED to either a control level approximating normal body temperature, a warm condition or two levels of cooling (**Fig. 1E**). We applied temperature manipulations using a block design of control-manipulation-control with transitions occurring at regular three-minute intervals (**Fig. 1E**). Manipulation temperatures were drawn at random and without replacement from the aforementioned set until its exhaustion, at which point the set was replenished and the sampling process resumed. We found that temperature near the tips of the probes tracked block changes, reaching asymptote within ~60 s of transitions (**Fig. 1E, Extended Data Fig. 1B, C**), and that temperature changes fell off to minimal levels within 6 mm of the probe tip (**Extended Data Fig. 1E, F**). This *in vivo* characterization of the implant (**Extended Data Fig. 1**) confirmed that temperature manipulations were largely localized to striatal tissue and that manipulation blocks of three-minute duration would allow for assessing effects of striatal temperature on neural activity and behavior.

**Figure 1.**
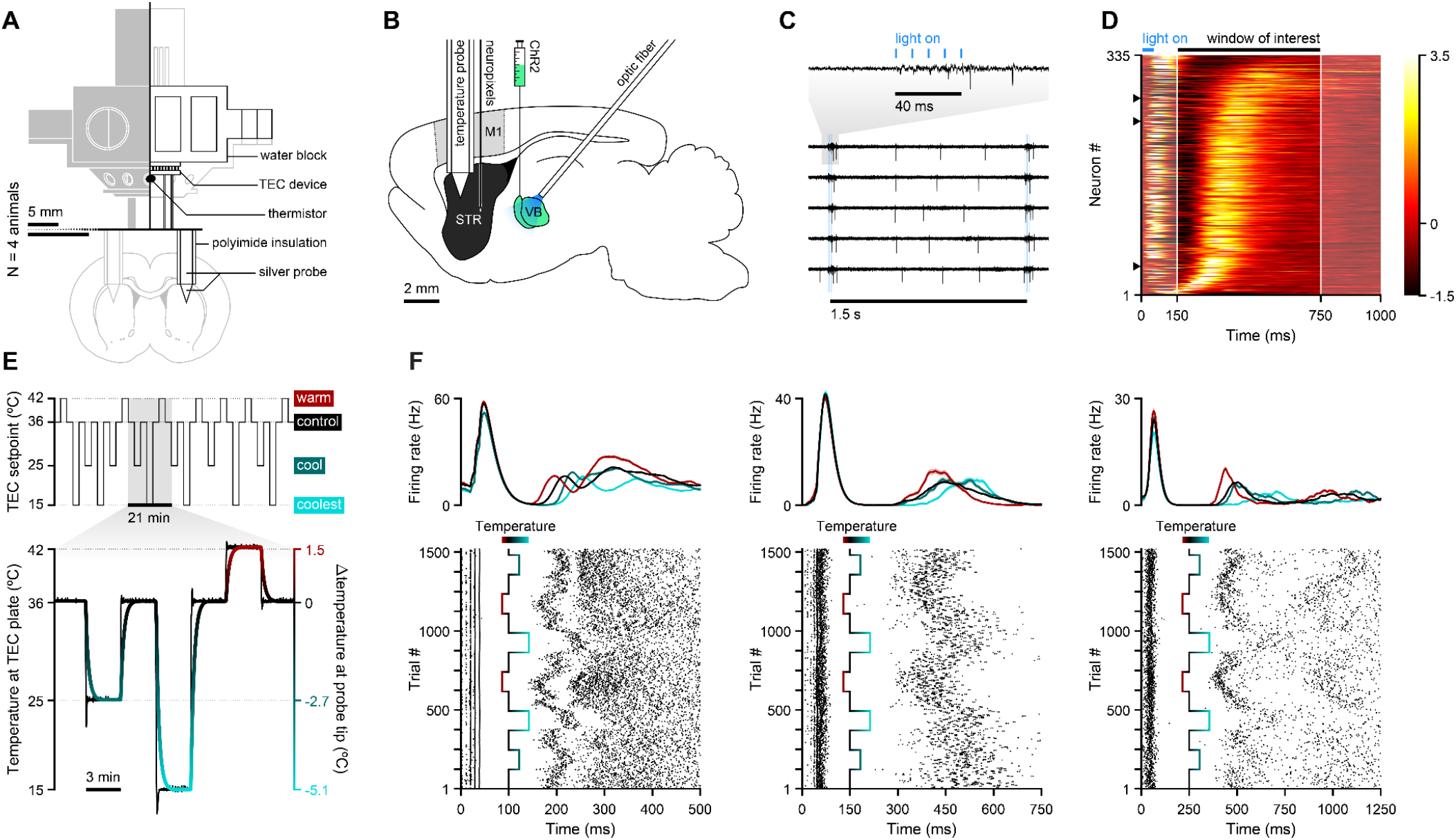
Temperature rescaled single striatal neuron responses monotonically in time. (**A**) Schematic of the thermoelectric device (TED). The thick horizontal line splits the diagram into two differently scaled subregions (see 5 mm scale bars). Top region is subdivided in the middle between a front view (to its left, with white strokes and gray fills) and a cutaway through the center of the implant. (**B**) Schematic of the preparation used to elicit, record and manipulate reproducible striatal dynamics under different temperatures. (**C**) VB thalamic stimulation protocol overlaid with five single-trial examples of evoked striatal voltage signals recorded in the control temperature condition. Blue ticks depict light pulses. (**D**) Normalized and smoothed peri-stimulus time histograms (PSTHs) of recorded striatal neurons (N = 335), under the control temperature. Units are ordered by their angular position in the subspace defined by the first 2 PCs describing firing dynamics between 150 ms and 750 ms from stimulation onset. This time window is delimited by white vertical lines and adjacent transparent patches. Arrowheads indicate example units shown in (F). (**E**) Top panel depicts the target setpoint of the peltier device over a representative session. Bottom panel depicts a representative segment of thermistor readings (black trace) at the lower plate of the TEC module shown in (A), illustrating the temperature manipulation protocol throughout experiments. Gradient-colored trace indicates the temperature as estimated from *in vivo* calibration experiments (Extended Data Fig. 1). (**F**) Activity of three putative striatal units aligned to the onset of VB stimulation. Top: Smoothed PSTHs split by temperature (mean ± s.e.m.). Bottom: Raster plots, wherein rows correspond to chronologically ordered trials and each tick mark to a unit-assigned action potential. The superimposed gradient-colored trace indicates temperature setpoint over the course of the recording session.

Next we characterized the features of neural activity that were affected by temperature, adapting a paradigm for optogenetically inducing patterns of striatal activity under anesthesia^13^. This approach allowed us to generate large numbers of highly reproducible bouts of population activity free from neural variability directly related to ongoing behavior, and was thus well-suited for studying the direct effects of temperature on striatal activity. Briefly, we used a viral strategy to express Channelrhodopsin-2 (ChR-2) in the ventrobasal complex (VB, **Fig. 1B**), a somatosensory thalamic area that projects directly to the striatum, and indirectly may influence striatal activity through thalamocortical projections. Three weeks post-infection, under urethane anesthesia, we implanted an optic fiber over VB, a single insulated silver probe of our TED into DS and an adjacent Neuropixels^34^ silicon probe (**Fig. 1B, Extended Data Fig. 2A, B**). We then delivered 50-ms trains of blue light pulses at 100 Hz once every 1.5 s and recorded DS neural activity (**Fig. 1C, D**). Stimulation caused a brief volley of striatal activity, followed shortly thereafter by reproducible patterns of firing across the population over hundreds of milliseconds after the last light pulse had been delivered (**Fig. 1D**). For assessing the effect of temperature on neural responses we focused on this longer lasting activity because, unlike the initial volley, later responses reflected autonomous dynamics of the system as it settles as opposed to the initial direct response to stimulation.

While the general patterning of individual-neuron responses over time was maintained across different temperatures, the time course of this pattern systematically varied depending on temperature, advancing more slowly the colder the temperature, and more quickly when temperature was raised above baseline (**Fig. 1F**). To quantify temperature-dependent warping in the time course of neural responses, we computed a scaling factor for each neuron in each of the four temperature conditions (see methods, **Extended Data Fig. 3A**). Temporal scaling as opposed to shifting of responses provided a significantly better explanation of the effect of temperature on firing rates across the population (**Extended Data Fig. 3**). Across all recorded neurons, distributions of response dilation were ordered inversely as a function of temperature (**Fig. 2A, B**), with the majority of cells exhibiting time-contracted firing profiles (dilation < 0%) under warming, and time-dilated firing profiles (dilation > 0%) under cooling conditions. At a population level this led to systematic differences in the speed with which population state advanced along its typical trajectory in principal component (PC) state space (**Fig. 2D**). Temperature also modified baseline firing rate of striatal neurons, however, this effect was distinct from the effect on temporal scaling. Whereas temperature produced a monotonic effect on neural temporal scaling across the sampled temperatures (**Fig. 1F, Fig. 2A, B, D**), baseline firing rates varied non-monotonically as a function of temperature, with both warming and cooling of striatal tissue relative to a physiologically normal control value resulting in lower firing rates (**Fig. 2C**).

**Figure 2.**
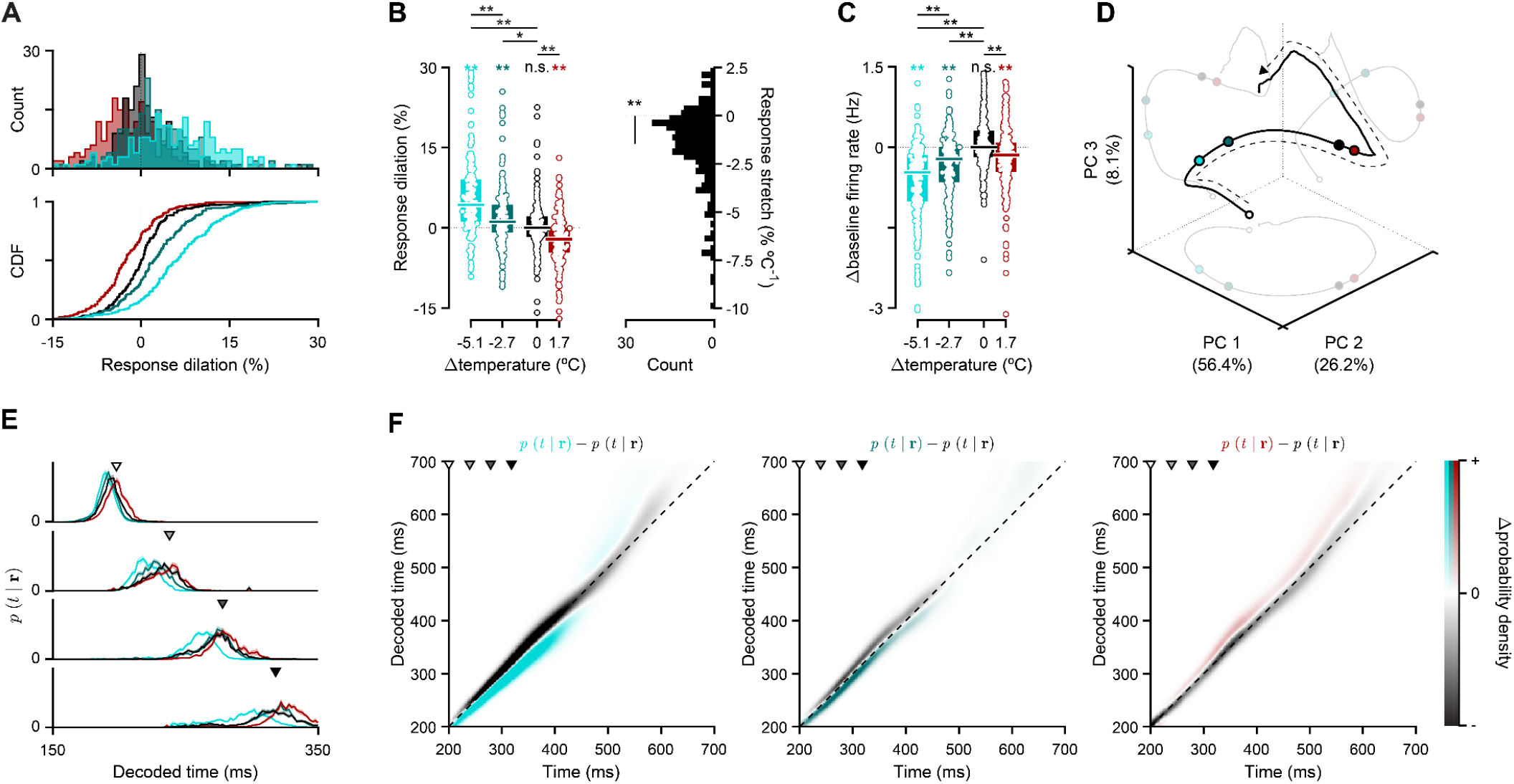
Temperature rescaled neural population activity in time leading to graded monotonic modulation of decoded time estimates. (**A**) Distribution of percentage change in temporal scale relative to control (dilation, N = 251, see methods). Histograms (top) and corresponding cumulative density functions (CDF, bottom) of dilation conditioned on manipulation temperature (same color scheme as in Fig. 1). (**B**) Left: Distributions of neuronal response dilations as a function of striatal temperature change. Markers represent temperature-split response dilations for each cell. Boxplots show population medians (horizontal thick lines) and i.q.r. (colored bars). Right: Distribution of stretch in neuronal responses. (**C**) Markers represent temperature-split change in baseline firing rate for each cell. Boxplots show population medians (horizontal thick lines) and i.q.r. (colored bars). (**D**) Average control population activity projected onto its first 3 PCs (solid black line), and temperature-split averages (colored markers) projected onto that control trajectory at the time point indicated by the last arrowhead in (E) and (F). Ghosted versions of these visualizations show their 2D projections onto all possible combinations of PCs 1, 2 and 3. The dashed black arrow indicates the direction of time. (**E**) Decoded posterior probability of time given the state of striatal populations at four arbitrary time points, averaged separately across trials for each temperature condition (mean ± s.e.m.). (**F**). Difference between trial-averaged decoded estimates for manipulation (left: extreme cooling; middle: mild cooling; right: warming) and control conditions, shown for the time window highlighted in Fig. 1D. Grayscale arrowheads indicate time points used in panel (E). The identity line is shown in dashed black.

Previous studies have shown that estimates of elapsed time decoded from striatal populations can predict timing judgments^22,23^. To assess whether temperature effects resembled endogenous, decision-related variability in population activity during behavior and its impact on the readout of decision variables, we decoded elapsed time from the population under different temperatures. Briefly, we first characterized the “typical” temporal profiles of striatal responses using a subset of control trials, and then applied a probabilistic decoding approach to estimate elapsed time based only on the observed state of the recorded population in remaining trials. Estimates of elapsed time derived from ongoing population activity systematically led ahead and lagged behind true time during warming and cooling blocks, respectively. This can be observed by sampling the output of the decoder at discrete delays from stimulation onset (**Fig. 2E**), and more continuously by subtracting decoder output during control blocks from that during the three different manipulation conditions (**Fig. 2F**). Relative to control, cooler temperatures gradually shifted decoded estimates of time earlier, and the warmer temperature shifted decoded estimates of time later. These data demonstrate that, under anesthesia, the temporal scaling of neural response profiles by temperature can reproduce features of endogenous variability in the time course of population activity and time encoding shown in previous studies to correlate with timing judgments.

### Striatal population speed predicts temporal judgments across tasks with different immobility requirements

A number of studies have observed that ongoing behavior may correlate with timing performance^35–40^. This raises the possibility that previously observed correlations between the speed of neural population activity and timing judgments might reflect the encoding of kinematic features by striatal populations^41,42^ combined with the adoption of embodied strategies for timing^43^. Such a scenario may indicate a more indirect role for striatal population dynamics in the decision process. To examine this possibility, we studied DS population dynamics (**Extended Data Fig. 2C, D**) during behavior in two versions of a temporal discrimination task (**Fig. 3A**) that differed in the degree to which animals were free to move around the behavioral box during interval estimation.

**Figure 3.**
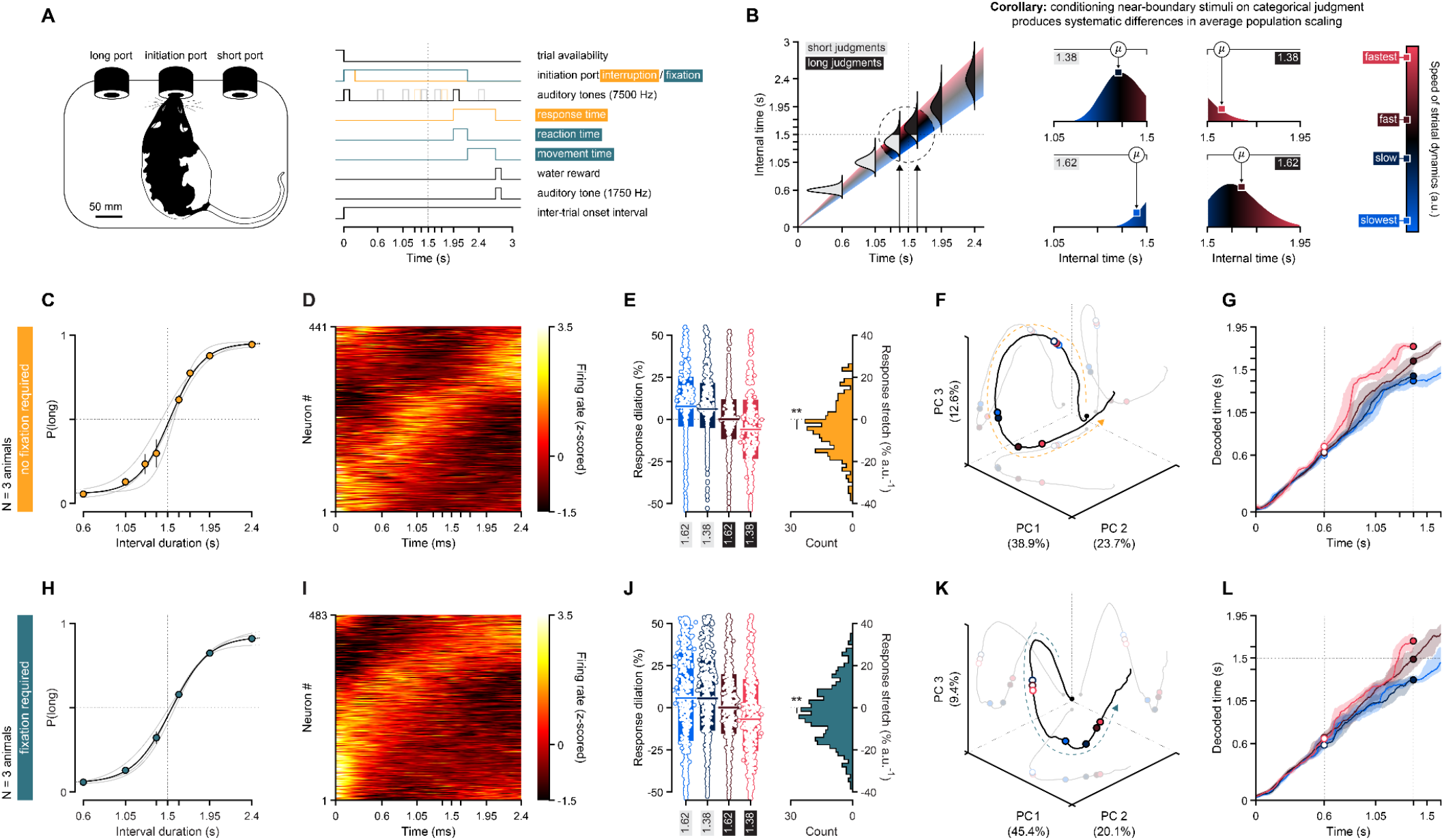
Conditioning neural activity on behavior produced monotonic modulation of population speed and decoded estimates of elapsed time. (**A**) Schematic of behavioral setup (left) and event diagram illustrating a correct trial (right) in both the *no fixation required* (orange) and *fixation required* (blue-green) versions of the interval discrimination task. (**B**) Left: Schematic of hypothetical relationship between trial to trial variability in the speed of striatal dynamics, blue indicating slower, and red indicating faster, and the resulting distributions of internal time estimates at each interval offset. Assuming a fixed internal decision boundary, long or short judgments are colored as black or gray portions of the distributions. Right: Schematic depicting hypothetical, graded shifts in average internal timing speeds when conditioning on choice for the two intervals nearest to the decision boundary. Specifically, short (long) stimuli incorrectly categorized as long (short) should reflect pronounced biases towards faster (slower) than usual dynamics, and correctly categorized short (long) stimuli should reflect milder biases towards slower (faster) dynamics. (**C**) Performance in the *no fixation required* version of the task (N = 3). (**D**) Normalized and smoothed PSTHs of all striatal neurons (N = 441) recorded during presentations of the longest stimulus in our set (2.4 s). Units are ordered by their angular position in the subspace defined by the first 2 PCs describing firing dynamics. (**E**) Left: Distributions of neuronal response dilations for each near boundary stimulus-choice pair introduced in (B). Markers represent response dilations for each cell and stimulus-choice condition. Boxplots show population medians (horizontal thick lines) and i.q.r. (colored bars). Right: Distribution of stretch in neuronal responses. (**F**) Average population activity from (D) projected onto its first 3 PCs (solid black line), and averages (colored markers) from each stimulus-choice pair projected onto that reference trajectory at stimulus offset for the short near-boundary stimulus (1.38 s). Ghosted versions of these visualizations show their 2D projections onto all possible combinations of PCs 1, 2 and 3. The dashed black arrow indicates the direction of time. (**G**) Average decoded M.A.P. estimates for each stimulus-choice condition. Colored solid lines and patches represent medians and i.q.r. across 512 concatenations (see methods), respectively. (**H**-**L**) Same as (C-G) for the *fixation required* task variant (N = 3 animals, N = 483 neurons).

We trained rats, in both versions of the task, to report intervals of time as either shorter or longer than a 1.5-s category boundary (**Fig. 3A**). Briefly, rats were placed in a rectangular behavioral box with three nose ports positioned at head level. Trials began with an auditory tone triggered by the subjects’ entry into the central “initiation” nose port. After an interval of silence during which subjects were either free to move about the box (*no fixation required*)^23^ or required to maintain their snout positioned in the central port (*fixation required*)^44^, a brief second tone was delivered. This second tone acted both as stimulus offset, defining the duration of the interval subjects were asked to judge, and a “go” cue, freeing subjects to report their choice at one of two equidistant ports on either side of the initiation port. Choices reported at one of the lateral nose ports following short stimuli (<1.5 s) and at the opposite lateral nose port after long stimuli (>1.5 s) were defined as “correct” and resulted in delivery of a water reward at the choice port. “Incorrect” and premature choices (in the no fixation required version of the task) or premature departures from the central port (in the fixation version), were punished with an error tone and a time penalty added to the inter-trial-onset interval.

Accuracy of timing judgments (**Fig. 3C, H**) and several key features of the recorded striatal population activity in relation to behavior were similar in the two paradigms. First, in both task variants, striatal population state continuously evolved through a set of non-repeating states during interval presentation (**Fig. 3D, I**). Furthermore, the time course of this evolution was systematically related to animals’ judgments, a relationship we examined using several approaches that facilitated comparison with the effect of temperature on neural activity described above.

Given previous observations that the time course of striatal responses covaries with duration judgments, conditioning trials of a given interval on animals’ judgments should result in distributions of neural population responses with systematically different temporal scaling (**Fig. 3B**). To test this prediction, we focused on intervals where long and short judgments were most balanced, corresponding to the two intervals nearest to 1.5 s. Because rats correctly categorized the majority of trials, even for these “difficult” stimuli, we expect the majority of any hypothetical distribution of population speeds across trials to result in population states at interval offset that lie on the “correct” side of the animal’s internal representation of the decision boundary. Thus, conditioning neural data on whether the animal judged 1.38 s as “long” or “short” should create two distributions of trials with respect to population speed: one with an average speed that is slightly slower than the overall average in the case of correct, “short” judgments (**Fig. 3B, right panel, top left**), and another with an average that is shifted to an even larger extent above the overall average toward faster speeds in the case of incorrect, “long” judgments (**Fig. 3B, right panel, top right**). Conversely, conditioning neural data on judgment of the 1.62-s interval should create one distribution of trials with an average speed that is slightly faster than the overall average in the case of correct, “long” judgments (**Fig. 3B, right panel, bottom right**), and another with an average that is shifted to an even larger extent below the overall average toward slower speeds in the case of incorrect, “short” judgments (**Fig. 3B, right panel, bottom left**). Using the temporal scaling metric applied to recordings under anesthesia above, we found that response dilation of individual neurons systematically varied in accordance with these predictions (**Fig. 3E, J**). As with manipulation of temperature, this led to systematic differences in the speed with which population state advanced along its typical trajectory in neural state space (**Fig. 3F, K**), and systematic differences in the speed of decoded time estimates derived from the population (**Fig. 3G, L**). Lastly, to gain a sense of how population speed might relate to choices across a broader range of stimuli, instead of conditioning neural activity on behavior, we conditioned behavior on the speed with which neural population activity evolved along its typical trajectory during the interval period in the two tasks. In both behavioral scenarios, rats were systematically biased toward making “short” judgments the more slowly the population state progressed on single trials, particularly for stimuli closer to the 1.5-s decision boundary, leading to a monotonic relationship between population speed and psychometric *threshold* (**Fig. 4A, Extended Data Fig. 5A, E**) - the stimulus duration at which the sigmoid crosses 50% probability of the animal making either a long or short choice. Thus, variability in timing judgments was associated with the same features of neural activity that were modified by temperature in the first set of experiments. Furthermore, this was true whether or not subjects were required to maintain their position in the initiation port during the interval stimuli, suggesting that decision-related variability in the time course of striatal responses did not result from encoding of detailed kinematic information alone. Next, having established that temperature provides a means of manipulating decision-related variability in striatal activity, we tested whether and how striatal temperature modified subjects’ decisions, movement parameters, or both.

**Figure 4.**
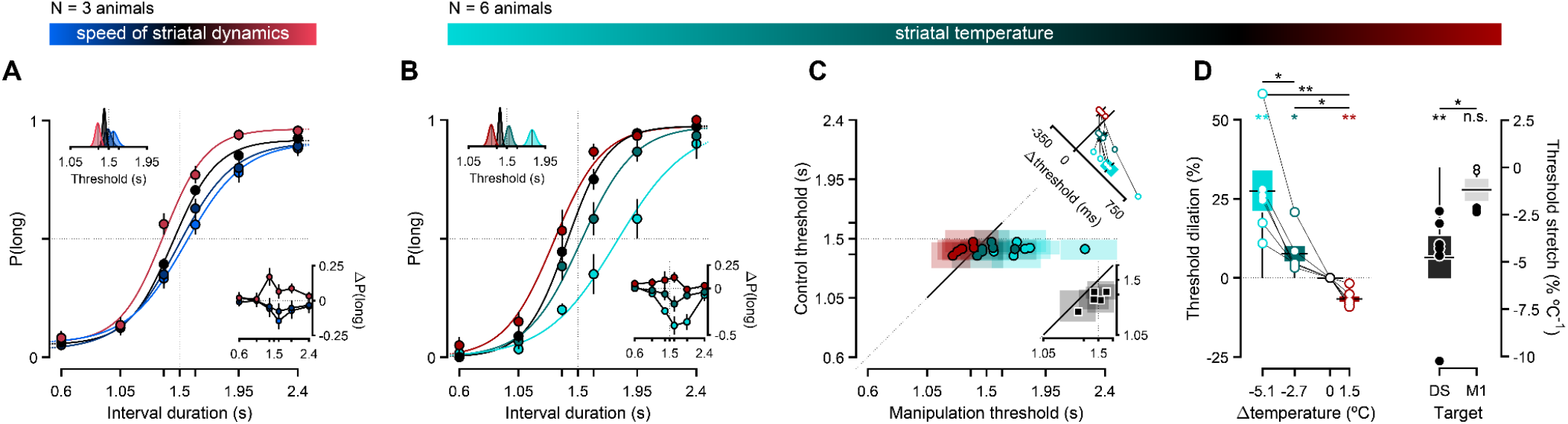
Conditioning behavior on population speed or temperature produced monotonic modulation of temporal judgments. (**A**) Performance in the *fixation required* version of the interval discrimination task conditioned on striatal population speed (N = 3, see methods): fast (red), typical (black), slow (light blue) or slowest (dark blue). Main axes: Psychometric curves fit to cross-animal averages of psychophysical data split by neural speed, respectively shown as solid lines and markers of matching color (mean ± s.e.m.). Bottom-right inset: Differences in proportion of long choices from each atypical speed condition to the typical speed condition (mean ± propagated s.e.m.). Top-left inset: Marginal posterior distributions of the threshold parameter for each speed condition’s psychometric fit. Solid black lines represent the maximum a posteriori (M.A.P.) point estimates. (**B**) Analogous to (A), but conditioned on striatal temperature (N = 6): warm (red), control (black), cool (teal) or coolest (cyan). (**C-D**) Effect of temperature on psychophysical thresholds. (**C**) Main axes: Markers represent M.A.P. estimates and transparent patches the corresponding 95% confidence intervals of threshold parameters fit to individual animals’ performance on control (y axis) versus manipulation blocks (x axis). Each animal contributes one data point of each color. Top-right inset: Distribution of threshold differences between manipulation and control conditions (mean ± s.e.m.). Bottom-right inset: Same as main axes, but with data pooled from 2 pilot experiments using a single cooling dose in either the *fixation* (N = 1) or the *no fixation* (N = 4) task variants (see methods and Extended Data Fig. 5). (**D**) Left: Distributions of threshold dilation as a function of induced temperature changes. Markers linked by solid black lines represent individual animals. Right: Distribution of threshold stretch split by manipulation target (DS: N = 6; M1: N = 4, see methods and Extended Data Fig. 6). Markers represent individual animals, and their size and color denote bootstrapped significance.

### Striatal temperature modified temporal judgments monotonically

We next implanted two additional groups of rats previously trained in either the version of the interval discrimination task not requiring (N = 4) or requiring fixation (N = 7), with a TED targeting its probe tips to DS (**Extended Data Fig. 2E, Extended Data Fig. 4D**). Critically, the no fixation required cohort and one rat in the fixation required cohort were implanted with a predecessor of the TED we described above, which was unsuited for prolonged extreme cooling and incable of warming. As such, for those animals temperature manipulations were restricted to a single mild cooling level.

In advance of temperature manipulations, and for both task variants, rats’ performance was virtually perfect for easy stimuli, progressively more variable as stimuli approached the categorical boundary and well described by a sigmoid psychometric function with a threshold close to the experimentally imposed decision boundary of 1.5 s (**Extended Data Fig. 4**). Strikingly, at the onset of temperature manipulations, all eleven subjects across the two task versions displayed a systematic shift towards short judgments when the striatum was cooled (**Fig. 4B-D**, Extended Data Fig. 5 B-D, F-H). Furthermore, subjects implanted with the latest TED version (**Fig. 1A**) exhibited bidirectional and monotonic changes in their discrimination behavior as a function of temperature: rats were more likely to report short judgments during cooling blocks, and long judgments during warm blocks (**Fig. 4B-D**), particularly for intervals nearer to the 1.5-s categorical boundary (**Fig. 4B**). Importantly, the larger the magnitude of the cooling manipulation, the larger the change in choice behavior. The systematic changes in the subjects’ judgments caused by temperature were most reliably captured by shifts in the threshold parameter of the psychometric function (**Fig. 4B, top left inset**). Thresholds tracked differences between control and manipulation temperatures in both sign and magnitude for all individual animals (**Fig. 4C, D**). Additionally, the circuit mechanism underlying the behavioral effects of temperature did not seem to involve overlying primary motor cortex (M1), through which the insulated portion of the probes passed (**Fig. 1A, B**), because direct manipulation of M1 temperature in an additional set of four rats (**Extended Data Fig. 2E**) performing the fixation version of the task produced significantly smaller effects on choice behavior (**Fig. 4D, right panel**) that did not reach significance (**Extended Data Fig. 6**) and where the time course of any trend appeared delayed relative to that of the effect on the DS cohort (**Extended Data Fig. 6D, Extended Data Fig. 9G**), consistent with volume conduction of cortical temperature manipulations down to the striatum (**Extended Data Fig. 1**). Thus, striatal temperature manipulations, shown above to produce changes in the population response under anesthesia that mimic decision-related temporal scaling of firing patterns, caused highly reproducible, parametric variation in a decision variable used by rats to guide duration judgments during the task. As in the recordings during behavior, this effect was robust to differences in task design with respect to the degree that animals were free to move during interval presentation. These data suggest that the systematic changes in timing judgments induced by temperature in the striatum are thus due to the effect of temperature on the temporal scaling of neuronal responses within populations of neurons that contribute to the decision.

### Striatal temperature modified movement kinematics non-monotonically

What can these results tell us about the nature of the functional involvement of striatal circuits in behavior? One of the most commonly observed effects of dysfunction in BG circuits is a change in the speed of movement, such as bradykinesia in the case of Parkinson’s disease. Indeed, inhibiting direct pathway striatal projection neurons has been shown to slow movement without affecting other aspects of its execution^44,45^. Since the anesthetized recordings described above revealed qualitatively distinct effects of striatal temperature on population speed (monotonic), and baseline activity levels (non-monotonic), we wondered whether there were any correlates of either or both effects to be found in measures of animals’ movements. We focused on fixation sessions, where both warming and cooling were applied, as bidirectional manipulation of temperature was observed under anesthesia to differentiate temperature’s effects on temporal scaling and baseline firing rate. We first tracked the position of the TED in video using a markerless tracking algorithm^46^. We then asked how temperature affected position over time either during the interval period (**Extended Data Fig. 7A-C**), or during choice movements (**Fig. 5A-C**).

**Figure 5.**
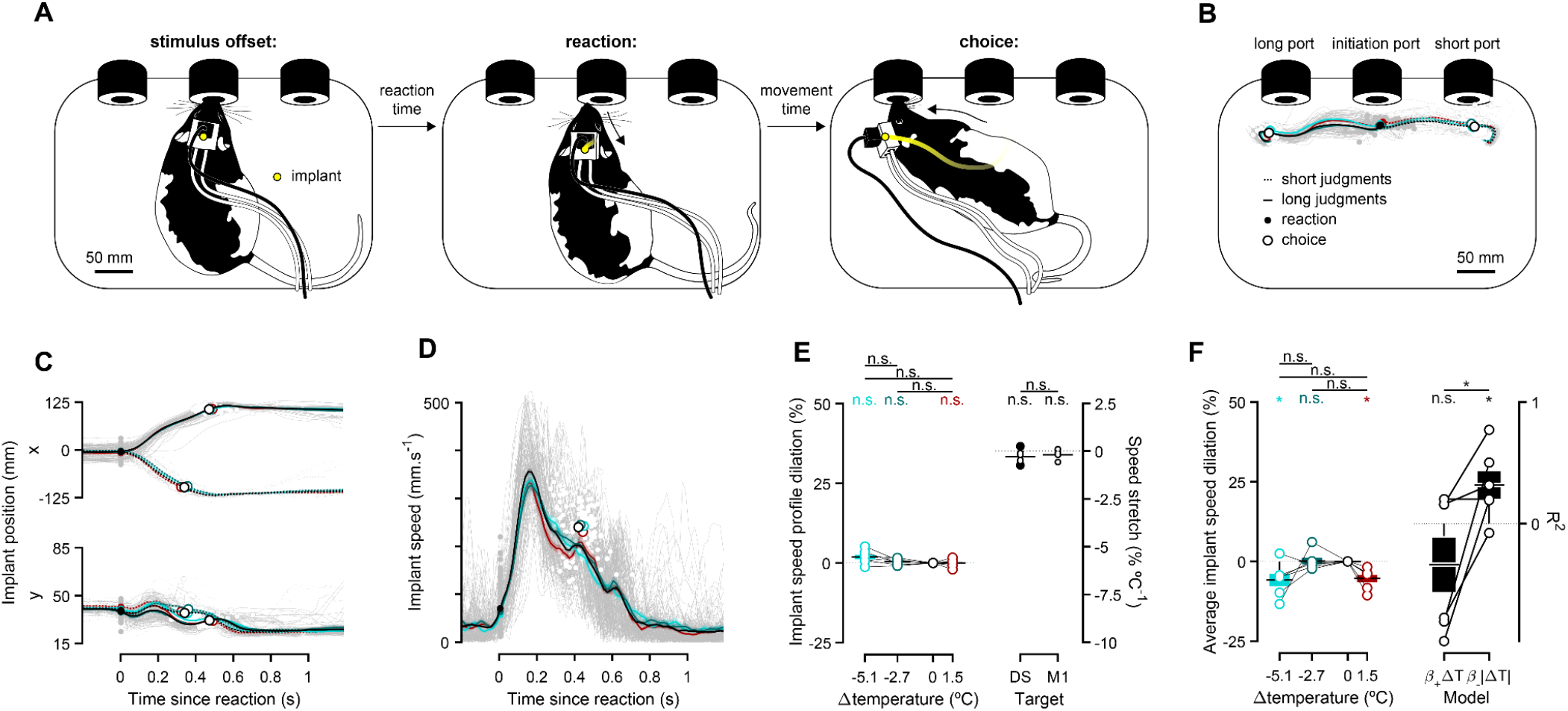
Temperature affected the kinematics of movement non-monotonically. (**A**) Illustration of 3 task epochs: stimulus offset (left), reaction (middle) and choice (right). (**B**) 2D trajectories of implant position aligned to reaction from a representative example animal. Dashed (solid) lines correspond to short (long) choices with individual control trials ghosted in the background, and condition-split averages on top. Filled and open markers show position at reaction and choice, respectively. (**C**) Same as in (B), split into horizontal (top) and vertical (bottom) coordinates. (**D**) Same data as in (C), combined into an overall speed metric. (**E**) Implant speed profile dilation for DS animals (left) and stretch for DS and M1 animals (right). (**F**) Left: same as in (E, left) for average implant speed dilation of DS animals. Right: Model comparison between 2 one-parameter models of temperature’s effect on average implant speed dilation in DS animals. The goodness of fit of the monotonic (left) and the non-monotonic (right) model, as measured by its coefficient of determination (R^2^), is shown for individual animals (open markers) and cohorts (mean ± s.e.m.). The *β* coefficient was constrained to only have positive (negative) values in the monotonic (non-monotonic) model. Color scheme indicates temperature as in previous figures.

We did not observe any significant monotonic effect of temperature in any case. Rats were positioned similarly during the interval period, and followed similar paths from the fixation port to the choice ports across the different temperatures (**Fig. 5B, C**). In addition, the speed profiles with which rats executed their choices were not scaled monotonically in time as a function of temperature (**Fig. 5D, E**). However, we did observe a modest non-monotonic effect in average movement speeds as a function of temperature (**Fig. 5F**), as evidenced by absolute temperature change being a better predictor of average speed over animals than signed temperature change (**Fig. 5F**). This was in stark contrast to the clearly monotonic effect of temperature on decisions and temporal scaling of population activity. Next we analyzed animals’ time to react to stimulus offset and the time taken to move to the choice port. We did not observe significant differences in either reaction times or movement times depending on temperature across rats (**Extended Data Fig. 7D-I**), however, we again observed a trend wherein both warming and cooling appeared to delay movement initiation following stimulus offset (**Extended Data Fig. 7F, J**), similar to the observed effects of temperature on both average movement speed (**Fig. 5F**) and baseline striatal firing under anesthesia (**Fig. 2C**). Accordingly, endogenous variability in baseline firing rates recorded in the fixation version of the interval discrimination task covaried with reaction times (**Extended Data Fig. 8**). These observations suggest that the temporal evolution of striatal population state controls the time course of decision variables used by rats to determine *when* to do *what* action, but that this feature of neural activity it is not what determines the moment by moment kinematics of movement execution (consistent with the observation that striatal lesions disrupt the sequencing of behavioral motifs, without affecting execution of motifs themselves^47,48^). Instead, we find evidence that the striatum may provide a lower dimensional gain signal coded in the firing rate of neurons that controls overall vigor^49–53^.

## DISCUSSION

Previous work has demonstrated that the speed of neural population activity along reproducible trajectories in neural space can correlate with variability in the timing of actions^4,54–56^ and in time-dependent decisions^6,23,57^. Here we show that experimental temperature manipulations in the striatum can be used to slow down or speed up both patterns of neural activity and the temporal evolution of latent decision variables that rats use to guide judgments of duration, an effect that was not explained as a temperature-dependent modulation of movement kinematics. We use the word *judgment* to encompass a range of potential processes related to the behavior, including both the continuous estimation of latent temporal decision variables, such as the time-varying value of particular actions, as well as ultimately the categorical choice of the animal. It has long been appreciated in fields as diverse as neuroscience, robotics, and artificial intelligence that control of movement is likely facilitated by a hierarchy of control mechanisms^58,59^. The observation that an intervention capable of modulating striatal population speed differentially impacts evolution of decision variables and those required for the timing of movement is consistent with the proposal that the BG act as a mid-level controller of movement, important for selecting among^9^, linking^12^, or modulating^60^ different actions but not so much the details of their execution. Indeed, lesioning dorso-lateral striatum in a motor timing task that involves timed transitions between elements of a motor sequence renders rats unable to perform the sequence^42^. Observations that kinematic information can be decoded from the striatum during tasks that involve timing^41,42^ are consistent with this view as well, because BG circuits likely benefit from access to continuous information about the state of commands that are sent to effector systems to guide when to produce particular actions, even if they are not responsible for sending detailed kinematic control signals to downstream circuits.

Such a hierarchical view is reminiscent of the observation that cooling an orofacial region of motor cortex in singing mice slows certain aspects of the song timing while leaving individual note durations unchanged^61^. Interestingly, tonic inhibition of the direct feedforward pathway of the BG at its initiation point in the striatum can produce a slowing of movement^45^, suggesting that the influence of the BG on control of parameters related to vigor may not be through dynamics in the higher dimensional space of population firing, but rather through a low dimensional, gain-like, modulation of motor programs that are largely implemented by circuitry elsewhere. Consistent with this we found that the non-monotonic effect of striatal temperature on baseline firing rates (**Fig. 2C**) resembled a pattern in rats’ behavior where both warmer and cooler temperatures produced delayed (**Extended Data Fig. 7F, J**) and slower choice movements (**Fig. 5F**).

While the effects of temperature on timing judgments were robust in the first sessions of manipulations (**Fig. 4**), these effects diminished with experience (**Extended Data Fig. 9A, D, G**). Though we had sought to minimize opportunities for animals to learn to adapt to the temperature manipulations by applying them in blocks of trials lasting only three minutes, always interspersed with a control temperature block (**Fig. 1E**), and alternating manipulation sessions with “washout” sessions where temperature was not manipulated (**Extended Data Fig. 4D**), we wondered whether animals might have learned to adapt to the effects of temperature. To assess this possibility, we performed a set of behavioral experiments in which the boundary between intervals rewarded for short judgments, and those rewarded for long judgments was shifted in the same blockwise manner (**Extended Data Fig. 9B**) as was temperature during the previously described experiments (**Fig. 1E**). This manipulation was devised to create a similar scenario to that created by striatal temperature manipulations in terms of unsignalled and surprising feedback that could be used by the animal to adapt their decision-making strategy to maximize rewards. Indeed, animals developed the ability to shift their decision thresholds over a small number of sessions, similar to the observed diminishing effects of temperature on temporal judgments (**Extended Data Fig. 9**). Thus, while temperature manipulations might also produce physiological adaptations that render neural systems more robust to future temperature variations^24^, we found that the changing impact of temperature on timing judgments was consistent with learning to compensate for decision variables whose features might be changing. These data suggest that in general the behavioral impact of manipulating neural systems should be evaluated continuously from the moment the manipulations begin, as opposed to evaluating end-point or average effects alone, as the adaptation ability of animals may be capable of overcoming what are initially significant effects on performance^62^.

During behavior, animals continuously interact with the environment, and this interaction drives significant amounts of neural activity across many brain areas. Pinpointing features of neural activity that underlie latent processes, such as aspects of cognition, in the face of behavior-driven neural activity is a difficult problem requiring multiple approaches. One approach involves studying neural activity across behavioral tasks that vary along dimensions that are orthogonal to the process of interest. For example, signals related to a decision and not its report should be invariant to movement conditions of the task^63^. In the current study, we found that variability in the speed of striatal population responses correlated with subjects’ temporal judgments across two task variants that differed in the degree that subjects were free to move about during stimulus presentation. *A priori*, it is impossible to determine whether such correlations reflect a causal relationship from that neural activity to behavior, an unobserved source of neural activity elsewhere that directly causes both striatal neural activity and behavior, or potentially behavior that causes striatal neural activity through reentrant sensory input. Previous analysis of high speed video during the non-fixation version of the temporal judgment task we study here demonstrated that decision-related information appeared in striatal activity several hundred milliseconds before it appeared in outward behavior, arguing against decision signals being driven by the sensory consequences of behavior alone^23^. Nonetheless, the existence of such varied possibilities underlies the need for so-called causal manipulations to help determine whether neural activity is a cause, corollary or a consequence of behavior. Thus, to characterize the impact of temperature on neural activity, we sought to remove behavioral sources of neural variability altogether^64^ by focusing on population dynamics elicited by optogenetic activation of VB thalamus under anesthesia. Importantly, in these experiments we did not seek to recapitulate the exact circuit mechanism by which population dynamics are produced during timing behavior. However, VB does integrate somatosensory signals that originate in areas throughout the body, including whisker and orofacial regions. These regions would receive mechanical stimulation at interval onset when rats insert their snout into the initiation port, and thus, VB thalamus represents one potential source of precisely timed^65^ information about the onset of interval stimuli.

What circuit mechanisms might give rise to task-relevant striatal population activity studied here? Previous views have suggested that the BG largely inherit their patterns of activity from inputs, often hypothesized to originate in cortex^4,66,67^. However, if time-varying striatal activity underlying temporal judgment were simply inherited from some other brain area, striatal temperature manipulations would be expected to produce a minimal shift and not a significant rescaling of behavior^28^. As in the vocal control circuit of songbirds, our data thus appear to be inconsistent with a mechanism where the relevant dynamics are simply inherited by the brain area targeted for temperature manipulations, in our case the dorsal striatum. In songbirds, there is evidence that a combination of local circuit mechanisms in pallial area HVC^68^ and a larger reentrant circuit involving HVC and multiple other brain areas are involved in generating the temporally patterned activity underlying song timing^26,28^. A similar scenario may underlie the mechanisms for generating temporally patterned striatal activity involved in temporal judgments. First, while most network modelling efforts that use neural dynamics for computation have relied at least in part on recurrent excitation, recent work suggests that it may be possible for a largely inhibitory, striatum-like circuit to produce complex spatio-temporal dynamics given sustained excitatory input^69^. The striatum may also represent one stage in a larger reentrant circuit involving multiple brain systems, where the larger circuit contributes to generation of dynamic patterns of activity that govern the evolution of decision variables. In this view, delays or advances induced by cooling or warming would accumulate with each cycle through the circuit, resulting in temporal rescaling with temperature. Such a circuit could in principle involve cortex, BG structures and thalamus^70^, or subcortical areas such as downstream BG structures, superior colliculus and thalamus^71^. However, our data suggest that any reentrant circuit mechanism involving cortex does not include primary motor cortex, as temperature manipulations there had negligible effects on choice behavior, consistent with previous studies demonstrating that manipulating motor cortex does not affect expert behavior in motor timing tasks^30,38^. However, orbito-frontal and medial frontal cortical areas have been shown to encode temporal information during both motor timing and temporal judgment tasks, albeit less accurately than the striatum^6,7^, and cooling of medial frontal cortical structures has been shown to delay movements^30^, potentially indicating involvement of frontal cortical structures. In addition, the activity of midbrain dopamine neurons correlates with and can directly cause changes in timing judgments^72^, suggesting that dopaminergic neuromodulation may additionally tune the time course of network activity through its action on striatal circuits.

It has been proposed that timing processes are distributed in the brain^2^, and that networks of neurons implicitly possess a rich capacity to act as timekeeping mechanisms through the time-varying patterns of activity they tend to produce during behavior, sometimes termed a “population clock”^3^. While the data presented here strongly support this hypothesis, in principle the kinds of computations performed by earlier more algorithmic, information processing accounts^73^ of timing might well be embedded in the type of population activity we describe here. This possibility is reflected in a growing belief that the brain performs many of its computations through dynamics^74^.

Our percepts, thoughts, and actions are continuously intertwined and regulated in time, and indeed understanding the neural basis of temporal processing has been argued to be a necessary prerequisite for general models of cognition^75^. Yet understanding how the brain appropriately orders and spaces information along the temporal dimension to encode signals that are functions of time has been an enduring challenge for neuroscience. Here we provide compelling evidence that the time course of activity in populations of striatal neurons directly influences the time course of a timing process used to guide decision-making. The data not only imply a causal link between temporal scaling of a population response and a temporal basis for computation in the brain, but argue that functions related to more abstract and low level aspects of behavior can, at least under some circumstances, be dissociated at the level of BG circuits. In the hierarchy of behavioral control, striatal dynamics appear to act at an interface between cognition and motor function to help guide *whether* and *when* to produce *what* actions, but not the details of movement execution. Understanding the precise circuit mechanisms responsible for establishing and modulating the timescale of neural activity in these circuits, and which specific computations this activity subserves, represent important future directions of inquiry if we are to understand how the brain draws on internally computed information to produce adaptive and intelligent behavior.

## METHODS

### Subjects

A total of 32 adult Long-Evans hooded rats (*Rattus norvegicus*) between the ages of 6 and 24 months were used in this study. 2 rats were used in an acute experiment aimed at characterizing the spatiotemporal profile of our temperature manipulation. Another 4 animals were used for an acute experiment combining electrophysiological recordings, temperature manipulation and optogenetic stimulation. 26 wild-type males were trained in the interval discrimination task (across the *fixation required* and *no fixation required* variants), of which 15 were chronically implanted with a custom TED that allowed for temperature manipulation experiments, 6 were implanted with 32-channel tungsten microwire moveable array bundles (Innovative Neurophysiollogy): 3 unilaterally (previously published data^23^) and 3 bilaterally. 5 were used in behavioral manipulation experiments. Prior to surgery, animals were kept in pairs in transparent cages with HEPA (High-Efficiency Particulate Air) filters on a 12-hour light-dark cycle (with lights ON at 8 am), at 21 °C and relative humidity of 50%. All experimental procedures were performed during the light phase of the cycle. Animals used in behavioral experiments had *ad libitum* access to food and were water-deprived. All experimental procedures were in accordance with the European Union Directive 2010/63/EU and approved by the Champalimaud Foundation Animal Welfare Body (Protocol Number: 2017/013) and the Portuguese Veterinary General Board (Direcção-Geral de Veterinária, project approval 0421/000/000/2018).

### Behavioral setup

The behavioral apparatus consisted of a 36 cm tall, 22.5 cm wide, and 35 cm long plastic storage box (TROFAST, Ikea) with three floor-level custom nose ports, a speaker (LS00532, Pro Signal) nearing the top of the opposite wall and a custom-made lid that provided uniform lighting and allowed for overhead video recordings (Flea3 FL3-U3-13S2, Point Grey Research Inc.) through an aperture. Each cylinder-shaped nose port was made up of 3D printed components housing a white light emitting diode (LED), an infrared (IR) emitter-sensor pair that enabled the detection of port entries and exits and the accompanying printed circuit board (PCB) (Champalimaud Foundation Scientific Hardware Platform). Additionally, the two lateral ports (positioned symmetrically around the central one) were each equipped with a metallic spout connected to a 20 mL water syringe via a solenoid valve (LHDA1231215H, Lee Company). All sensors, actuators and peripherals were respectively monitored, controlled and kept in the same temporal reference frame using a custom finite state machine implemented by a microcontroller I/O board (Arduino Mega 2560, Arduino) and an interfacing PCB (Champalimaud Foundation Scientific Hardware Platform). Finally, detected port events and other task-relevant behavioral data were timestamped, serially communicated to a Windows 10 desktop computer and stored as a parseable text file using a custom python script. Video was acquired at 60 FPS with a resolution of 1280 × 960 pixels in 8-bit grayscale using Bonsai^76^.

### Behavioral training

Leading up to the experimental sessions reported in this paper, animals were first trained in 2 hour-long daily sessions 5 times a week in various “tasks” of increasing complexity. During this stage, which we termed *Poking101*, rats were progressively introduced to the following rules: (un)lit ports are (un)responsive, meaning that nose-poking into a lit port will cause it to turn off and trigger some task event, whereas doing so at an unlit port is inconsequential; entering a lit lateral port results in a reward delivery of 25 uL of water paired with a brief auditory tone (1750 Hz, 150 ms); entering the central port when it is illuminated initiates a trial and *can* lead to both lateral ports lighting up. This is contingent on the animal’s snout continuing to interrupt the IR beam at the center port for the entirety of a “fixation delay” (*fixation required* version), or the animal not making a premature entry at either lateral port during the “stimulus presentation delay” (*no fixation required* version). Whichever the task variant, this imposed delay starts off at 0 s and is adaptively marched up towards 3 s (within and across sessions) and consists of an interval of silence demarcated by two brief auditory tones (7500 Hz, 150 ms). Failure to withhold premature departures from the central port (*fixation* version) or choices (*no fixation* version) causes the current trial to be aborted, eliciting an error tone (150 ms of white noise) and adding a timeout of 15 s to the already ticking 9-s inter-trial-onset interval (ITOI). Once animals were able to reliably maintain fixation at the central port (*fixation* version) or defer choices (*no fixation* version) for 3 s, training on the interval discrimination task began^23,37^. In it, instead of waiting for a fixed amount of time and collecting a reward at either lateral port once it elapsed, rats were asked to wait for a variable delay on each trial and to then categorize it as either shorter or longer than a boundary of 1.5 s. “Short” judgments were registered at one of the lateral nose ports and “long” judgments at the opposite one. Rewards were contingent on stimulus and judgment categories being the same. When this was not the case, an error tone (150 ms of white noise) was played and a time penalty of 10 s was added to the ITOI. Pairs of stimuli symmetric about the categorical boundary were gradually introduced (from easiest to hardest) into the discrete sampling set that animals experienced, until reaching *I* = {0.6, 1.05, 1.38, 1.62, 1.95, 2.4} s (*fixation* version) or *I* = {0.6, 1.05, 1.26, 1.38, 1.62, 1.74, 1.95, 2.4} s (*no fixation* version). A correction-loop procedure was used such that, following 3 incorrect categorizations of any given stimulus, only that stimulus was presented to the animal until its associated error count dropped below 3. This training mechanism was disabled during manipulation sessions. It took ~3 months for rats to reach asymptotic performance.

### Thermoelectric Device (TED)

#### Design

We used a custom-made implantable TED (weighing ~30g) based on the Peltier effect to manipulate temperature in neural tissue. The implant consisted of a heat dissipation module, a thermoelectric cooling (TEC) module (01801-9A30-12CN, Custom Thermoelectrics), a 10kΩ thermistor (9707204, Farnell) and two 15-mm long sharpened silver probes. These were insulated down to, but excluding, the tips with a thin layer of PTFE low density thread seal tape (00686081520745, Gasoila) and polyimide tubing. The main distinguishing factors between the implant’s initial prototype (used in the single cooling dose during the *no fixation required* interval discrimination task variant) and its final version (used on the DS and M1 bidirectional manipulation cohorts performing in the *fixation required* task variant), were that the former was constructed with a passive aluminum heatsink (ICKS25X25X18,5, Fischer Elektronik) and 0.5-mm thick probes insulated with 1-mm wide polyimide tubing (95820-11, Cole-Parmer), whereas the latter had active heat dissipation via a water block (WBA-1.00-0.49-AL-01, Custom Thermoelectrics) and 1-mm thick probes insulated with 2-mm wide polyimide tubing (95820-13, Cole-Parmer). This was used in tandem with a peristaltic pump (200-SMA-150-050, Williamson), male and female Luer adapters (WZ-45504-00, Cole-Palmer) and the required interfacing tubing (WZ-06407-71, Cole-Palmer), allowing for a continuous flow (~15 mL/min) of room temperature water through the water block’s inner chambers. The TEC’s upper plate was glued to the bottom of the heatsink using thermal glue (TBS20S, TBS), which was also used to secure the thermistor at the center of this module’s lower plate. Finally, the two sharpened silver probes were soldered onto the TEC’s lower plate (one on each side of the thermistor) using a mixture of lead (419424, Farnell) and silver solder (SDR-9703-030, Custom Thermoelectrics), at a distance of 5 mm from each other. This inter-probe spacing corresponds to two times the ML stereotaxic coordinate of all our DS-targeted implants. Lastly, an RJ45 (85513-5014, Molex) connector was added on top of the heat sink and a custom 3D-printed spacer was mounted on its bottom, both secured using epoxy resin (2022-1, Araldite).

#### Closed-loop control

The implant was plugged into a custom-made PCB (developed by the Champalimaud Foundation Scientific Hardware Platform and available upon request) via an ethernet cable. This PCB implemented a proportional-integrative-derivative (PID) controller that was designed to bring the implant’s thermistor measurement to any experimenter-defined target temperature (within the TEC’s range of operation). Briefly, the thermistor readout was continuously compared to the current temperature setpoint in order to compute an absolute error term (*proportional* channel), a cumulative error (*integrative* channel) and an instantaneous change in error (*derivative* channel). These three error terms were then linearly combined, with weights set by the resistive and capacitive components of the hardware modules that implemented them, and used to modulate the control current driving the TEC. This negative feedback mechanism was optimized so that the target temperature could be reached with negligible delays, steady-state errors and over/undershoots. The resulting closed loop control allowed for stable, safe and transient temperature manipulations, as it required less user intervention, monitorization and arbitration than open loop alternatives. The PID’s setpoint was communicated through a serial communication pin from an additional Arduino Mega 2560 board that implemented the temperature manipulation protocol *per se*, meaning it controlled both when to transition into a new block and which temperature to transition to. All block types lasted for 3 minutes, except for control ones in our single cooling dose experiment, which were twice as long to accommodate slower heat dissipation due to this initial experiment’s characteristic passive heatsink. In all cases, block transition times and target temperatures were respectively signalled via a brief digital pulse and an additional serial communication port to the task-implementing Arduino board. Both the PID-implementing PCB and the block-controlling Arduino were connected to a computer running Windows 10, where a LabView-based (National Instruments) graphical user interface (TEC visualizer, Champalimaud Foundation Scientific Hardware Platform) enabled online visualization and saving of digitized thermistor temperature measurements (sampled at 100 Hz). Lastly, to prevent irreversible tissue damage in the eventuality of a partial compromise of the closed loop system leading to its “opening”, an additional failsafe mechanism was implemented in the PCB’s firmware, ensuring that the TEC was automatically disabled if the registered thermistor temperature ever dipped below 0 °C or rose above 55 °C.

#### Calibration

A calibration curve between different set temperatures at the lower plate of the TEC module and temperature measurements at the tip of the silver probes was derived from an acute preparation with an anesthetized rat. Lower plate temperature was set to each value in *T =* {5, 15, 20, 25, 30, 45} °C, in blocks of 4 minutes, always preceded and followed by a control block of the same duration (*T =* 36 °C). Temperature at one of the tips of the implant’s probe was measured by a second thermistor glued along the probe axis to the polyimide insulation layer. In a separate acute experiment, we positioned a thermistor probe angled at 30° at different distances to the implant’s tip (*D* = {0.3, 0.5, 0.9, 2, 5.75} mm), and for each of them repeated the aforementioned blocked calibration procedure (but with manipulation temperatures drawn from the set *T =* {15, 25, 42, 45} °C). Lastly, all implants were tested individually post-assembly to ensure their respective TEC modules were functioning steadily and properly calibrated, using the same 4-min block protocol in warmed agarose gel (1.5%), which has similar thermal properties to brain tissue^77^.

### Surgical procedures

#### Acute temperature measurements and calibration

Rats were anesthetized with 2.0-4.5% isoflurane. Animals’ body temperature was continuously monitored and maintained at 35 °C by a rectal probe connected to a closed-loop heating system (FHC, https://www.fh-co.com). After being anesthetized and before making the first incision, we administered dexamethasone (2 mg/Kg), carprofen (5 mg/Kg) and a saline solution of atropine (0.05 mg/Kg) subcutaneously. We stereotaxically targeted the DS unilaterally (+0.84 mm AP, +2.5 mm ML from Bregma ^78^. Immediately following these procedures, animals were perfused for histological confirmation of the measurements’ location.

#### Viral injections

Following the same procedure for induction and maintenance of anesthesia used for acute temperature measurements and calibration, we stereotaxically targeted the ventrobasal complex (VB) of the thalamus for viral delivery (−2.3 mm AP, ±2.8 mm ML, 6.6 mm DV from Bregma^78^). We injected 300nL of rAAV5-CamKII-hChR2(H134R)-EYFP (titer ~10^12 GC%; University of Pennsylvania Vector Core) using an automated microprocessor controlled microinjection pipette with micropipettes pulled from borosilicate capillaries (Nanoject II, Drummond Scientific). Injections were performed at 0.2 Hz with 2.3 nL injection volumes per pulse. For all injections, the micropipette was kept at the injection site 10 minutes before withdrawal. Craniotomies were then covered with Kwik-Cast (WPI) and the skin was closed with sutures (Vicryl, Ethicon Inc.). Animals were allowed to fully recover on a warming pad and returned to the home cage when fully alert. During the 3 days following surgery, animals were given carprofen (5 mg/Kg, SC).

#### Acute optogenetic stimulation, extracellular recordings, and temperature manipulation

Following 3-6 weeks for viral expression, 4 rats were anesthetized with two doses of urethane, the first at 0.7 g/Kg of body weight and the second at 0.35 g/Kg 20 minutes after. Additionally, we administered dexamethasone (2 mg/Kg), carprofen (5 mg/Kg) and a saline solution of atropine (0.05 mg/Kg) subcutaneously. Animals were then kept with isoflurane at 0.5-1% until at least 30 minutes before electrophysiological recordings began. Animals’ body temperature was continuously monitored and maintained at 35 °C by a rectal probe connected to a closed-loop heating system (FHC, https://www.fh-co.com) throughout the experiments. We opened a large rectangular craniotomy over the left hemisphere (4 mm AP by 3 mm ML from Bregma^78^), centered in the same target location as the chronic implants. A 300 µm diameter and 0.37NA optic fiber (Doric) was targeted to VB (−2.3 mm AP, ±2.8 mm ML, 6.2 mm DV from Bregma^78^), inserted at a 39° angle and secured with blue light cured self adhesive resin cement (RelyX(tm) Unicem 2 Self-Adhesive Resin Cement, 3M). A small silver ground wire was inserted under the skull of the opposite hemisphere. A TED similar to the one used for chronic implants (with a single silver probe at a 90° angle relative to the heat sink, to accommodate the geometrical demands of the experimental preparation) was lowered to the same DS target location (+0.84 mm AP, −2.5 mm ML, 4 mm DV from Bregma^78^). This modified device was calibrated and behaved similarly to the ones used for chronic manipulations. Finally, a Neuropixels probe (Phase 3A Option 3, IMEC ^34^) was placed caudally relative to the temperature probe, and slowly lowered to target (5 to 6.5 mm DV) and allowed to stabilize in the tissue for at least 30 minutes before starting recordings and stimulation protocols. Seldomly, and for longer recording protocols, an additional dose of urethane was necessary to maintain anesthesia (0.2 g/Kg). An 473 nm LED source (Doric) was connected to the implanted optical fiber using a patch cord (400 μm core, 0.48 NA) and set to 3.5-5.5 mW at the end of the fiber and controlled using a dedicated arduino that was also responsible for switching the block temperature identity through a serial communication with the TEC controller. Each stimulation trial consisted of a single train of 5, 1-ms long, pulses at 100 Hz (each train lasting 50 ms in total). Each trial was separated by a period of 1.5 seconds. Electrophysiological and peripheral synchronization (LED and temperature probe) data were simultaneously acquired using SpikeGLX software (https://billkarsh.github.io/SpikeGLX/) at 30kHz. Local-field potential gain and action potential gain were set at 250 and 500, respectively, and split at 300 Hz. Each block at a specific temperature lasted 3 minutes. Temperature identities were drawn, without replacement, from the available set of 3 temperatures and were always intercalated with a control block. This protocol was repeated twice for a total of 2 blocks for each manipulation condition. Immediately following these procedures, animals were perfused for histological confirmation of the measurements’ location.

#### Chronic TED implants

Rats (N = 15; 5 in the single cooling dose pilot experiment, 6 and 4 in the bidirectional striatal and M1 temperature manipulation experiments, respectively) underwent surgery around 3 months after they started training. During the implantation of the TED rats were anesthetized with 2.0-4.5% isoflurane. Animals’ body temperature was continuously monitored and maintained at 35 °C by a rectal probe connected to a closed-loop heating system (FHC, https://www.fh-co.com). After being anesthetized and before making the first incision, we administered dexamethasone (2 mg/Kg), carprofen (5 mg/Kg) and a saline solution of atropine (0.05 mg/Kg) subcutaneously. We stereotaxically targeted the DS bilaterally (+0.84 mm AP, ±2.5 mm ML from Bregma^78^). Two craniotomies and durotomies matching the diameter of the silver probes were made. 5 support screws were placed: 1 in the occipital plate, 2 posterior and 2 anterior to the location of the craniotomies. The cranial bone was covered with self-curing dental adhesive resin cement (Super-Bond, C&B) to improve adherence to the dental acrylic used to secure the implant. The TED was then slowly lowered perpendicular to the brain surface to a depth of 4 mm from cortical surface. The craniotomies were covered with Kwik-Cast (WPI) and the implant was fitted into place and secured with several layers of dental acrylic (the first of which mixed with gentamicin). The procedure ended with suturing (Vicryl, Ethicon Inc.) the skin anterior and posterior to the implant. Animals were allowed to fully recover on a warming pad and returned to the home cage once fully alert. Animals were then individually housed to minimize implant damage. During the 3 days following surgery, animals were injected once a day with carprofen (5 mg/Kg, SC). Animals were allowed to recover for a week after the surgery with food and water *ad libitum*.

#### Chronic electrophysiology implants

These procedures followed a protocol similar to the one used for the implantation of the chronic TED implants. For a detailed description, see^23^.

### Temperature manipulation protocol

Following one week of recovery since surgery, all animals were again water deprived and gradually resumed behavioral training. Once they were performing at approximately pre-surgical levels in the interval discrimination task described above, they were subjected to temperature manipulation sessions. These 2-h sessions were divided in 3-minute blocks: control blocks, in which the TED was set to body temperature (~36 °C), always interleaved with manipulation blocks, in which the TED was either set to 25°C in the single cooling dose pilot experiment, or one of 3 manipulation doses (15, 25 and 42 °C) in all bidirectional manipulation experiments. Manipulation temperatures were drawn at random and without replacement from the aforementioned set until its exhaustion, at which point the set was replenished and the sampling process resumed. Sessions invariably started and ended with a control block and animals were not explicitly cued to block transitions. Manipulation sessions were interleaved with washout sessions, in which the TED’s controller was disabled, and correction-loop training was reinstated.

### Implant placement confirmation

Rats were sacrificed with transcardiac perfusion with phosphate-buffered saline (PBS), followed by 4% (wt/vol) paraformaldehyde (PFA). Following perfusion, brains were left in 4% PFA for 24 h and then moved to a 30% sucrose solution (wt/vol) in PBS for 2 to 3 days. For chronic electrophysiology, single-cooling dose and acute experiments, a vibratome was used to section the brain into 50 μm coronal or 40 μm sagittal slices, respectively. Coronal slices were stained with NISSL and sagittal slices series were alternated with NISSL or immunostained with a primary antibody against GFP (A-6455, Invitrogen) and a secondary antibody conjugated with AlexaFluor 488 (ab150077), and finally, incubated in DAPI. Images were acquired with a stereoscope (Lumar V12, Zeiss) or a slide scanner (Axio Scan Z1, Zeiss). For animals subjected to bidirectional chronic temperature manipulations, a 1 T MR scanner (ICON, Brucker) was used to collect MRI data. A T2-weighted structural image of the brains was collected using a Rapid Imaging with Refocused Echoes (RARE) pulse sequence. The sequence used had a repetition time of 2800 ms, echo time of 90 ms and a RARE factor of 12. The field of view was set to 28 × 15 × 20 mm^2^, the spatial resolution of the images was 150 × 150 × 150 μm^3^ or 80 × 80 × 80 μm^3^ and a matrix of 187 × 100 × 133 voxels was acquired after 8 averages during a 7-hour scanning.

### Data analysis

Unless otherwise stated, all data were analyzed using custom MATLAB 2019b (https://www.mathworks.com) scripts.

### Psychophysical data analysis

#### Preprocessing

Trials with reaction times greater than 1 s, or movement times greater than 3 s were labelled as outliers and excluded from all reported analyses. This resulted in less than 5% of all trials being removed. In order to make balanced comparisons across animals and temperature conditions, data from the initial 2 manipulation sessions of every chronically implanted animal were pooled together chronologically up to the point where there were 10 trials per stimulus for each manipulation condition and 40 trials per stimulus for the control temperature condition. The same pooling procedure was applied in reverse for the last 2 temperature and boundary manipulation sessions.

#### Psychometric function

We used the *Psignifit*^*79*^ toolbox to fit the following 4-parameter psychometric function to all interval discrimination data:

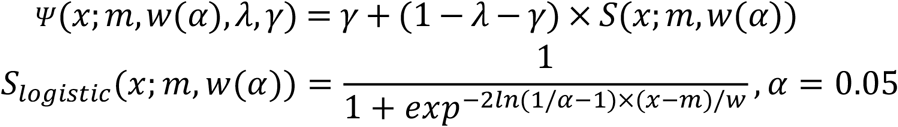

In this parameterization, a strictly monotonic sigmoid function *S* from the stimulus level *x* onto the unit interval [0,1], is specified by *m* = *S*^−1^(0.5) and *w* = *S*^−1^(1 − *α*) − *S*^−1^(*α*), namely the *threshold* and *width* parameters. This is independent of the choice of *S*, which in our case is the logistic function. The hyper-parameter *α*, which sets the span of *w* along the y axis, was set to 0.05. To account for stimulus-independent choices, *S*is scaled by two additional free parameters, *λ* and *γ*, which respectively control the upper and lower asymptotes of the psychometric function *Ψ*. The *λ* and *γ* parameters were fixed across temperatures at values found through fitting the corresponding control temperature data.

#### Dilation & stretch metrics

We adopted the dilation and stretch definitions from^28^. Briefly, *dilation* (*D*) of any scalar metric *x* (e.g., threshold M.A.P.), was calculated as the percent difference from unity in the ratio of a given temperature’s estimate over that of control.

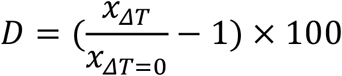

*Stretch* (*S*) was defined as the slope coefficient in a least squares linear regression using dilation as the response variable and the magnitude of our temperature manipulation (induced temperature differences around the implant’s tip) as the sole predictor.

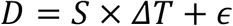

### Electrophysiological data analysis

Unless otherwise stated, what follows applies to all three electrophysiology data sets analysed in this paper: acute DS recordings (N = 394 neurons, across 4 animals), chronic DS recordings during the *no fixation required* (N = 441 neurons, across 3 animals), and the *fixation required* (N = 483, across 3 animals) versions of the interval discrimination task. We use the generic terms *condition* to refer to striatal temperature in the acute data set, and to stimulus-choice pairs in the two chronic data sets. The *reference* condition corresponds either to control temperature (acute) or to correct categorizations of the 2.4-s stimulus (chronic). Lastly, trial onset refers to the onset of VB stimulation (acute) or stimulus onset (chronic).

#### Preprocessing

In the case of the acute recordings, we focused on contiguous periods of stable activity (without abrupt changes in firing profile/rate), which in practice meant trimming the very beginning and/or end sessions where needed. For the 3 animals we recorded from chronically and unilaterally in the DS during performance in the *no fixation required* version of the interval discrimination task, preprocessing was done as described when this data set was first published^23^. For the remaining animals, we used a semi-automated offline approach to detect and sort recorded action potentials into well-isolated units and clusters of multi-unit activity. Detection, sorting and inference of the relative depth of each unit, were done using KiloSort2 (github.com/MouseLand/Kilosort2), whereas curation of the resulting clusters was performed using Phy (github.com/cortex-lab/phy). Prior to any of the analyses shown in the main figures, we further selected validated units with an intersectional approach that used firing rate and stability, as well as recording depth in the case of the acutely recorded data. Briefly, in order to survive this selection step, units had to: have a mean firing rate of 0.5 Hz or higher; be “stable” throughout the recording session, which we enforced by discarding units for which the Pearson correlation coefficient between average spike density functions computed with two random non-overlapping permutations of reference trials was lower than a 0.75 threshold; and in the case of the acute recordings, have been recorded at a contact that was later inferred to be in the DS, by comparing its position along the Neuropixels probe to a dip in the distribution of recorded cell depths, likely corresponding to a characteristic silence when transitioning from gray (cortex) to white (corpus callosum) to gray (striatum) matter.

#### Single neuron responses

Spike density functions were built on a trial-by-trial basis by first counting spike times in 2-ms bins and then convolving the resulting histogram with a causal kernel specified by a gamma distribution with shape and scale parameters of *k* = 2 and *θ* = 75 ms, respectively. Baseline firing rates in the acute recordings were computed in a 500 ms window preceding VB stimulation. To compute temporal scaling factors for each unit-condition pair, we first upsampled reference spike density functions by a factor of 10 and then warped them in time using 1000 scale factors, linearly spaced in the range [0.625, 1.75]. Both the upsampling and time-warping steps were performed using linear interpolation. Next, we linearly regressed all time-warped templates against the spike density function of each condition and stored the corresponding coefficient of determination (R^2^). The scale factor that maximized this template-matching metric, is what we operationally defined as the *temporal scaling factor* for that unit and experimental condition. In the case of control scaling factors, we split data into two random non-overlapping sets of trials and arbitrated which one was used to construct templates and which one was the target. Regarding response dilation and stretch, we used the same definitions from the psychophysical analysis section, except that scaling factors (*f*) replaced ratios of temperature over control estimates when calculating dilation:

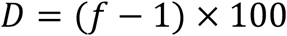

#### Low dimensional representations of population state

We used Principal Component Analysis (PCA) to enable visualization of striatal population trajectories in representative 3-dimensional subspaces. Briefly, we first averaged spike density functions across trials of a given condition and concatenated them into *N* × (*T* × *K*) matrices, where *N* is the number of neurons for that data set, *T* the number of 2-ms time bins relative to trial onset, and *K* the number of experimental conditions. After normalizing these data to have zero mean and unit standard deviation along the temporal dimension, we used trial-averages corresponding to the reference condition to find the three orthogonal directions that maximally captured variance in said data. Lastly, each condition’s trial-averaged population activity was then projected onto the subspace defined by these principal components (PCs), with reference trajectories plotted for all time points, and non-reference trajectories plotted only for arbitrary time points and projected onto the corresponding reference trajectory so as to prevent visual clutter.

#### Psychometric curves split by population state at stimulus offset

This analysis was adapted from its original introduction in^23^, and was only applied to the two chronic data sets in the current paper. Briefly, for all individual trials in each session with 5 or more simultaneously recorded units, we projected population activity at stimulus offset onto the median trajectory traversed by that striatal ensemble during the entire stimulus presentation period. We then normalized these projections by the length of that median trajectory. In addition to this temporal scaling metric, we computed an outlierness metric as the average point-by-point minimum distance between each trial’s trajectory and its session’s median trajectory (the 5% most extreme trials in these “outlierness” distributions were removed from all subsequent analyses). After pooling all normalized projections over all sessions and animals, we then partitioned, for each stimulus, the resulting distributions into *Q* groups. For ease of comparison, *Q* was chosen to match the number of conditions in the temperature manipulation experiments during behavior in the *no fixation required* (*Q* = *K* = 2) and the *fixation required* (*Q* = *K* = 4) variants of the interval discrimination task. Psychometric curves were then fit to trials from each group as described above.

#### Decoding time from ongoing population activity

We used a naïve Bayes decoder (flat prior) to continuously compute probability distributions over elapsed time using multi-session concatenations of putative striatal population activity aligned to trial onset.

Briefly, we:

1) Discretized time *t* into *B* 2-ms bins, such that *b* ∈ [1, *B*] and *t*_*b*_ ∈ [0, *T*]ms, with *T* = 1500ms for the acute recordings and *T* = 2400ms for the chronic recordings;

2) Fit an encoding model to each neuron *n* ∈ [1, *N*] at each point in true time *t*_*b*_:

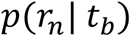

Which we determined empirically by querying cell-specific single-trial spike density functions *r*_*n*_ at the true time interval [*t*_*b*_, *t*_*b*+1_], and smoothing the resulting rate histograms with a gaussian kernel (*μ* = 0, *σ* = 10Hz). This was done using a subset of trials making up half of all reference trials;

3) Made conditional independence assumptions about neurons, regardless of whether or not they were simultaneously recorded:

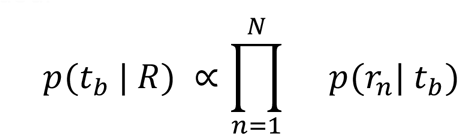

Where *R* = (*r*_1_, *r*_2_, …, *r*_*N*_) is a novel, to-be-decoded instance of concatenated population activity recorded at a known condition and point in true time since trial onset;

Used Bayes’ rule in creating a decoding model that linearly combined all individual neuron encoding models with a uniform prior over decoded time 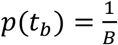:

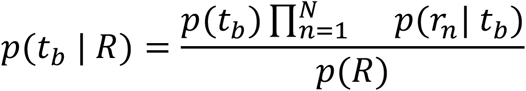

Where the probability *p*(*R*) for the population vector *R* to occur does not have to be estimated explicitly, as it indirectly follows from normalizing the posterior distribution *p*(*t*_*b*_ | *R*) such that it sums to 1 over all possible decoder outcomes, i.e., elapsed time as decoded from striatal ongoing activity.

Once the time-dependence in the responses of striatal cells recorded during a set of training trials is known, this Bayesian approach directly addresses the inverse problem: given the firing rates of the same cells, now recorded during previously unseen test trials, how likely is it for any and all *b* units of time to have passed.

### Continuous behavioral data analysis

#### Preprocessing

Full session 2-h videos recorded during the fixation version of the task were first cut into 12-s long clips (one per trial) aligned on stimulus onset. Offline tracking of the position of several implant features was performed using DeepLabCut^46^. Contiguous low confidence position estimates (likelihood < 0.85) of 6 samples or less were linearly interpolated and tracking time series were subsequently smoothed with a 6-sample median filter. In order to standardize units of distance across animals and sessions, we computed session-wise background frames by taking the median pixel intensities from a random sample of 3000 frames from each session’s full video. Next, we arbitrarily picked one background frame from a particular session of a particular animal as a reference, and computed the affine transformations that would best align each session’s background frame to that reference. These transforms were then applied to both the horizontal and vertical coordinates of each tracked feature and finally, converted to approximately SI units of distance by exploiting knowledge of the behavioral box’s dimensions and assuming no nonlinear optical distortions.

#### Implant speed

Instantaneous implant *speed* (*s*) was computed by taking the one-step forward difference of the average (across three implant features) displacement (*r*) with respect to sampling time (Δ*t*).

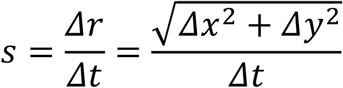

To compute speed profile dilation and stretch, we used the same template-matching approach described above for single-neuron responses, but using average condition-split speed across time instead of condition-split spike density functions. Lastly, average speed dilation was calculated in the same way as threshold dilation, with each condition’s average - the mean across single trial median speeds computed between reaction (the first detected exit time from the initiation port following stimulus offset) and 350 ms (~the time animals took to settle into a nose port) after choice (the first detected entry time at a choice port) - replacing threshold point estimates.

### Statistics

Unless otherwise stated, we used one-sample two-tailed t-tests whenever assessing the statistical significance of shifts in distributions, which we visually afforded with vertical solid black lines connecting the distribution’s mean to zero. When examining differences across distributions, we used either two-sample two-tailed t-tests when comparing striatal and motor-cortical stretch distributions, or repeated measures ANOVA followed by post-hoc contrasts with Tukey correction for multiple comparisons when comparing experimental dilation distributions across conditions but within cohorts. We visually afforded these two- and paired-sample tests with horizontal solid black lines connecting the two underlying distributions, offset in y for clarity. In all cases, we denote test outcomes near the respective visual affordance with the following notation: *, p<0.05; **, p<0.01; n.s., not significant. In the case of single animal stretch estimates, we assessed their statistical significance at a 5% level by bootstrapping. Specifically, we computed these point estimates for 1000 random samples per manipulation condition, constructed by sampling equal numbers of trials with replacement from the control condition while preserving stimulus identity. For each iteration, we then performed linear regression on bootstrapped dilations and stored the respective slope coefficient as that iteration’s stretch. Bootstrapped significance was consistently denoted by larger dark-filled markers, as opposed to smaller white ones.

## ACKNOWLEDGEMENTS

We thank Bassam Atallah and Caroline Haimerl for comments on versions of the manuscript and the entire Paton lab, past and present, for feedback during the course of this project. We would also like to thank the ABBE Facility and the Scientific Hardware, Histopathology and Rodent Champalimaud Research Platforms for unparalleled technical assistance. We thank Francisca Fernandes and Daniel Nunes for acquiring the MRI scans and Mauricio Toro and Renato Sousa for help with animal training. This work was developed with the support from the research infrastructure Congento, co-financed by Lisboa Regional Operational Programme (Lisboa2020), under the PORTUGAL 2020 Partnership Agreement, through the European Regional Development Fund (ERDF) and Fundação para a Ciência e Tecnologia (FCT, Portugal) under the project LISBOA-01-0145-FEDER-022170. The work was funded by an HHMI International Research Scholar Award to JJP (#55008745), a European Research Council Consolidator grant (#DYCOCIRC - REP-772339- 1) to JJP, a Bial bursary for scientific research to JJP (#193/2016), internal support from the Champalimaud Foundation, and PhD fellowships from FCT to FSR (SFRH/BD/130037/2017), BFC (PD/BD/105945/2014) and AIG (PD/BD/128291/2017). We thank the support of NVIDIA Corporation with the donation of the Titan X Pascal GPU used for this research.

## Author contributions

TM, FSR, MP, JJP devised the experiments, TM, FSR, MP performed all experiments, analyzed the data and drafted and edited the manuscript. BFC helped design and perform electrophysiology experiments and reviewed the manuscript. AIG performed a subset of temperature manipulation experiments during behavior and reviewed the manuscript. PERO devised and assisted in implementing the method of optogenetically stimulating reproducible striatal dynamics and reviewed the manuscript. JJP supervised all aspects of the project and drafted and edited the manuscript.

## Materials & Correspondence

The data and analysis code that support the findings of this study are available from the corresponding author upon reasonable request.

## EXTENDED DATA FIGURES

**Extended Data Figure 1.**
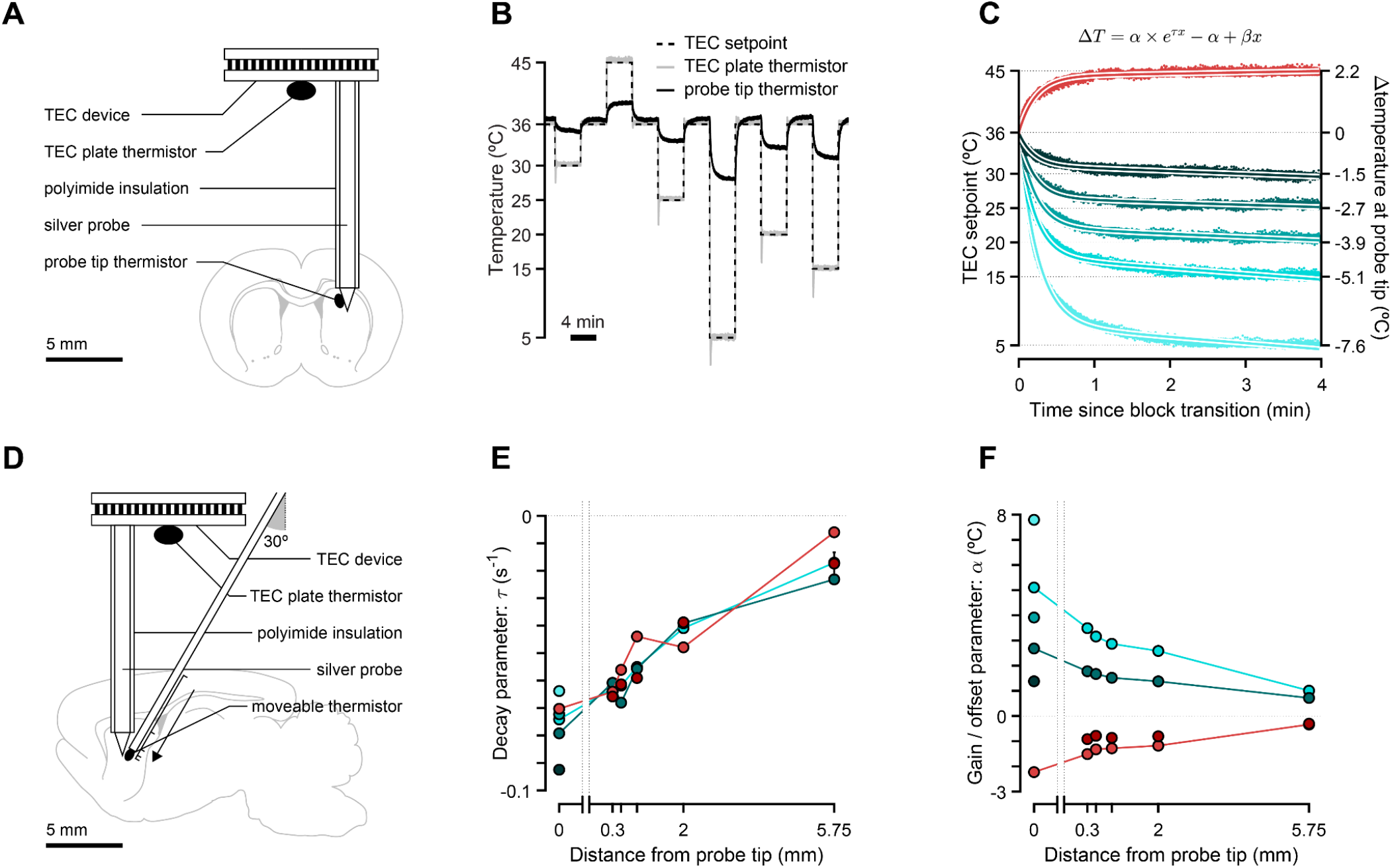
Spatiotemporal characterization of thermoelectric device (TED). (**A**) Schematic of the preparation in which we set our TED to one of several manipulation temperatures (*T* = {5, 15, 20, 25, 30, 45} °C) while measuring temperature at its lower plate and probe tip simultaneously. (**B**) Temperature measured at the TEC plate and probe tip thermistors. (**C**) Temperature traces measured at the probe tip thermistor during manipulation blocks aligned to block transitions. Solid lines represent model fits. (**D**) Schematic of the preparation in which we set our TED to one of several manipulation temperatures (*T* = {15, 25, 42, 45} °C) while measuring temperature at its lower plate a movable temperature probe simultaneously. (**E**) Decay parameters for models fit to manipulation temperatures (as shown in (C)) across the 2 experiments (manipulation temperatures that are common to both experiments are connected with solid colored lines). (**F**) Same as (E), but for the gain / offset parameter across all model fits.

**Extended Data Figure 2.**
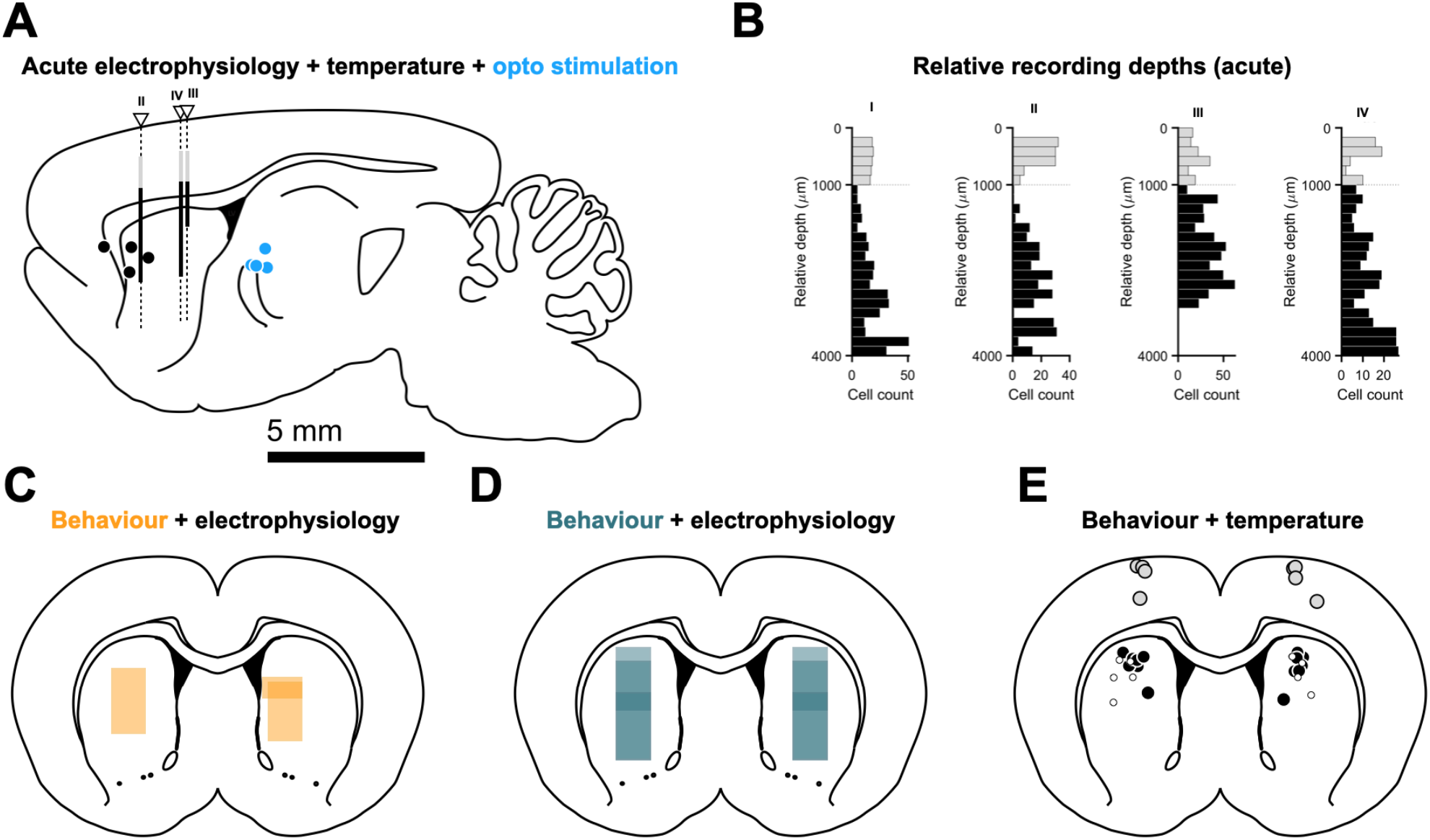
Histological reconstruction of TED, optical fiber and recordings probe placements for acute and chronic experiments. (**A**) Intermediate medial-lateral (ML) locations of TED probes (black markers), optical fibers (blue markers) and Neuropixels probes (white triangles) projected onto a reference sagittal slice (ML = 2.62 mm from Bregma). (**B**) Distributions of relative recording depths for all animals (N = 4) and recorded units (N = 335). Horizontal dashed line represents corpus callosum. Putative motor cortical and striatal neurons in gray and black, respectively. Histograms’ relative depth is overlaid in (A) using the same color scheme. We were unable to clearly identify the Neuropixels tract for animal I. (**C**-**E**) Intermediate anterior posterior (AP) location of microwire recording bundles in the *no fixation required* (C, orange, N = 3 animals implanted unilaterally), *fixation required* (D, petrol blue, N = 3 animals implanted bilaterally) and TED (E) probes for striatal (black markers, N = 6) and cortical (gray markers, N = 4) targets projected onto target coronal slice (AP = +0.84 mm from Bregma). White markers show implant locations for the *no fixation required* striatal cooling experiment (N = 5).

**Extended Data Figure 3.**
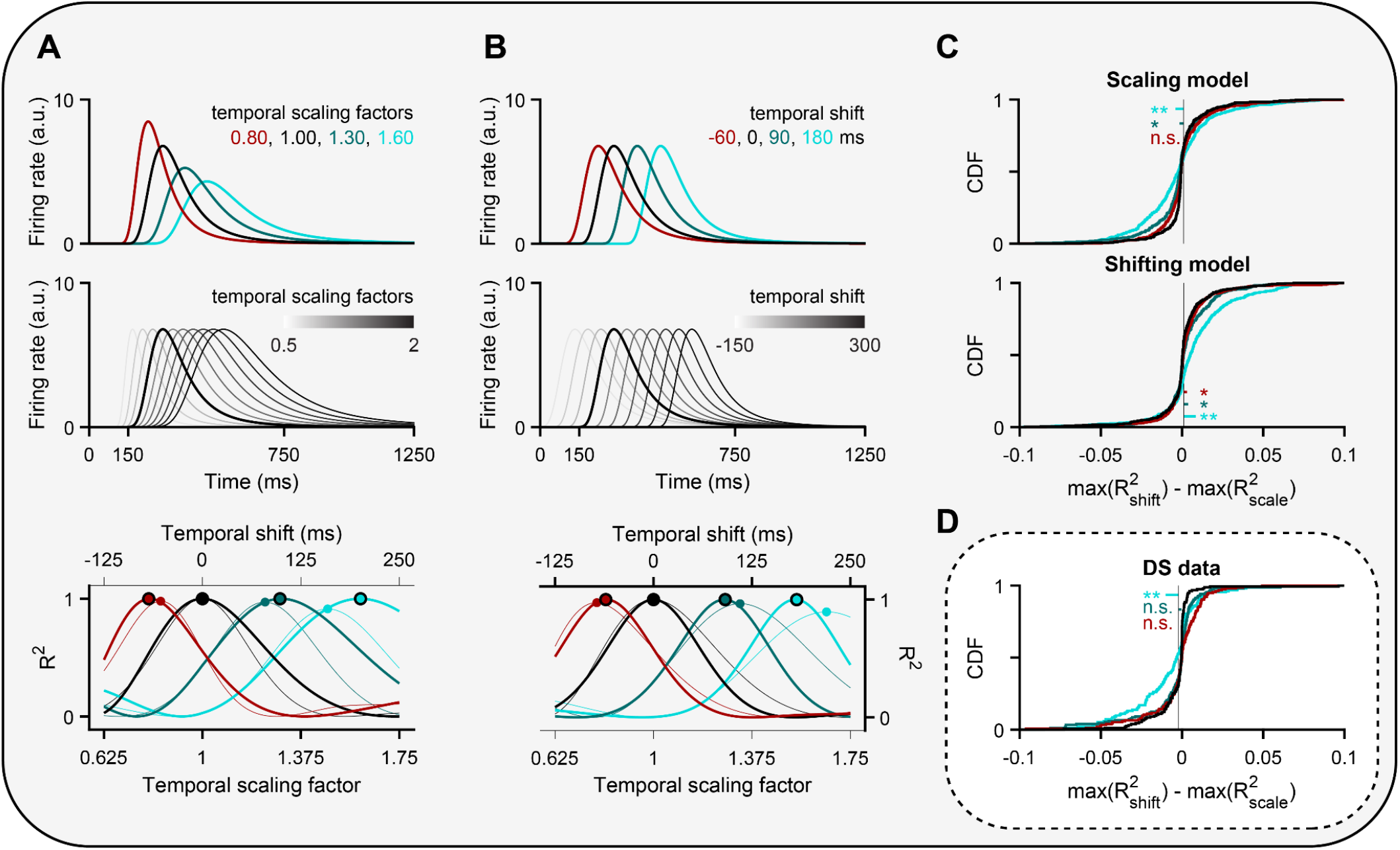
Temporal scaling as opposed to shifting provided a better account of temperature’s effect on neural activity. (**A**) Top: Simulated spike density functions exhibiting bidirectional and dose-dependent temporal scaling with temperature. Middle: Templates built by warping a control spike density function (thicker black line) in time by scale factors ranging from 0.625 (maximum contraction) to 1.75 (maximum dilation). Note that when applying this method to data, this control response is not the same as the one shown in the top panel, as the two are built using two non-overlapping random sets of control trials. Bottom: Thick lines represent the coefficient of determination (R^2^) for all scaled templates in the middle panel regressed against each of the target spike density functions shown at the top. We computed this objective function for each neuron-temperature condition pair and took its global maximum as the corresponding temporal scaling factor, highlighted here by the larger markers. Thinner lines and smaller markers depict R^2^ values for a similar regression procedure applied to a series of shifted, as opposed to scaled, templates (see B). (**B**) Same as (A), except that for artificially temporally shifted responses relative to control (top), temporally shifted templates (middle), and their regression outcomes (bottom). The thinner lines and smaller markers respectively represent the R^2^ curves and maxima resulting from regressing the scaled templates from A, middle against the shifted targets in B, top. Conversely, the result of regressing shifted templates against scaled targets is plotted in the same manner in (A, bottom). (**C**) To assess whether the effects of temperature on individual striatal responses were better accounted for by temporal scaling or shifting, we built two separate spiking models in which we either injected one effect or the other. Briefly, we modeled 500 control firing rate functions as gaussian bumps defined over 1.5 s with means spanning the interval from 150 ms to 750 ms (**Fig. 1D**) and a standard deviation of 50 ms. The amplitudes of the resulting probability density functions were rescaled so that their distribution of mean firing rates matched that of striatal data. Next, we created one additional rate function per neuron per manipulation condition by either shifting or scaling its control response in time. Again, the distribution of generative temporal scaling factors and shifts used was informed by the empirical distributions of these metrics extracted from striatal data. We then generated 150 spike trains of each condition per neuron by sampling spike times from inhomogeneous Poisson point processes with the aforementioned condition-specific responses as their time-dependent rate parameters. From this point on, we proceeded to analyze the resulting spike “data” in the exact same way we did for the striatal data, by first averaging trials within condition, generating libraries of templates and then computing temporal scaling factors and shifts. Finally, for each “neuron”-condition pair within each model, we stored the R^2^ values corresponding to the best-matching scaled and shifted templates and subtracted the former from the latter to build the distributions shown here at the top (scaling model) and middle (shifting model) panels. Thick solid sigmoidal lines represent the CDFs of each condition’s R^2^ difference. Thin vertical black lines denote control mean differences. Small horizontal colored lines link the respective means of the corresponding manipulation and control distributions. (**D**) Same as (C), but for striatal data.

**Extended Data Figure 4.**
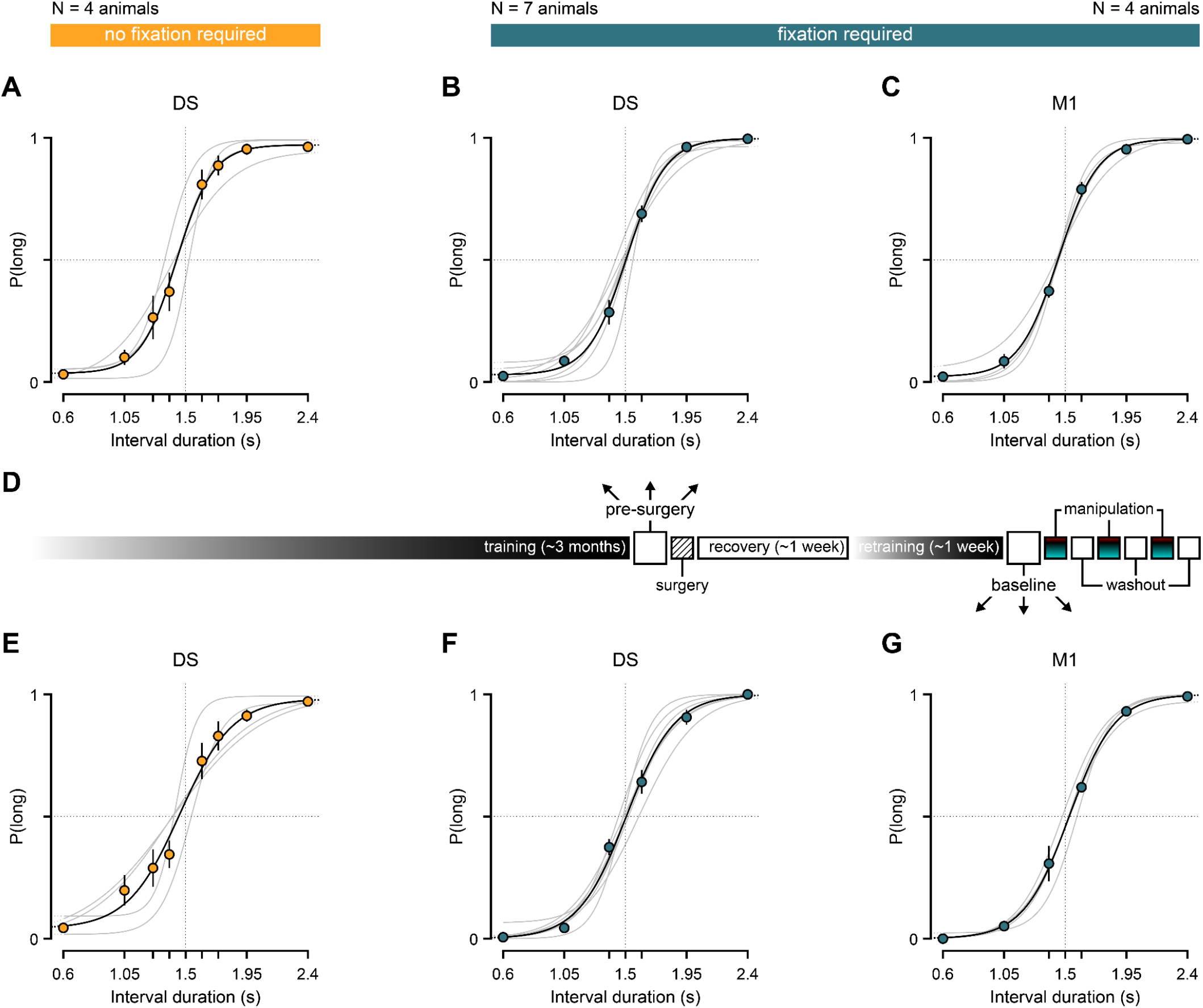
Discrimination performance in both task variants was qualitatively similar before and after TED implantation. (**A**) Discrimination performance of the cohort of rats that were trained on the *no fixation required* variant of the interval discrimination task and eventually implanted with the initial version of our custom TED targeting the DS (N = 4) on the last day of training before surgery. Gray lines are psychometric fits to individual animals. The black one is a fit to the average across animals. The underlying choice data is shown for the cohort average (mean +-s.e.m.). (**B**) Same as (A), but for the cohort of rats that were trained on the *fixation required* task variant and eventually implanted with the initial (N = 1) or final version of our custom TED targeting the DS (N = 6). (**C**) Same as (B), but for the cohort of rats that were eventually implanted with the final version of our custom TED targeting M1 (N = 4). (**D**) Schematized timeline for a typical experimental rat. Elongated bars illustrate long periods of time, whereas squares represent individual daily sessions, which, with the exception of surgeries, lasted for 2 hours. (**E**-**G**) Same as (A-C), but for the last post-surgery training session in advance of starting temperature manipulation sessions.

**Extended Data Figure 5.**
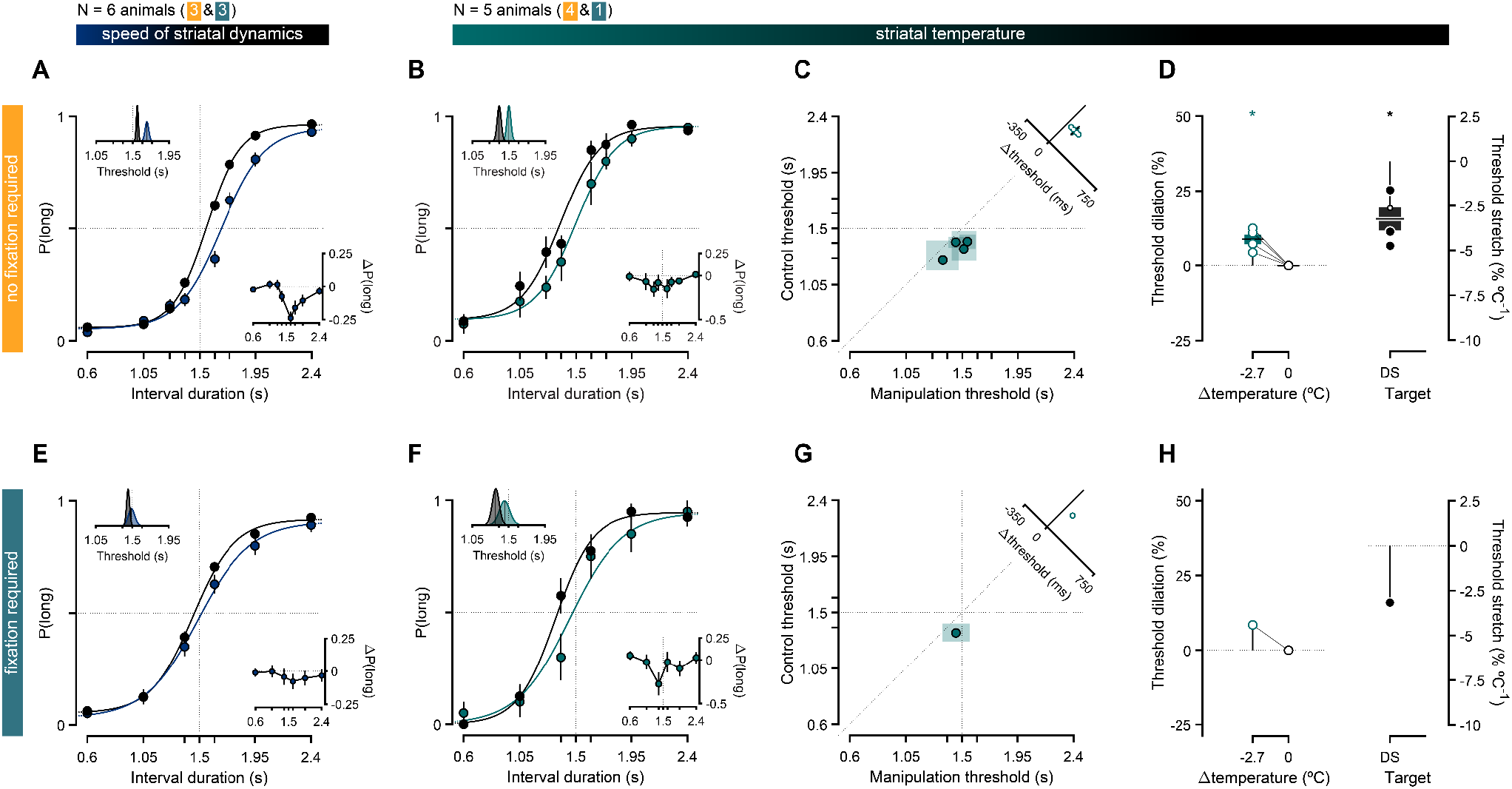
TEDs capable of a single mild cooling temperature produced qualitatively similar effects on timing judgments in both variants of the interval discrimination task. (**A**) Performance in the *no fixation required* version of the interval discrimination task conditioned on neural population speed (N = 3). Main axes: psychometric curves split by whether activity progressed more slowly (blue) or at a typical speed (black, see methods). Bottom-right inset: Differences in proportion of long choices from the slow speed condition to the typical speed condition (mean ± propagated s.e.m.). Top-left inset: Marginal posterior distributions of the threshold parameter for each speed condition’s psychometric fit. Solid black lines represent the maximum a posteriori (M.A.P.) point estimates. (**B**) Analogous to (A), but conditioned on whether striatal temperature was set to control (black) or a mild cooling (teal) dose (N = 4). Main axes: psychometric functions fit to cross-animal averages of temperature-split psychophysical data, respectively shown as solid lines and markers of matching color (mean ± s.e.m.). Bottom-right inset: Average differences in proportion of long choices from the mild cooling condition to control (mean ± propagated s.e.m.). Top-left inset: Marginal posterior distributions of the threshold parameter for each condition’s psychometric fit. Solid black lines represent the maximum a posteriori (M.A.P.) point estimates. (**C-D**) Effect of temperature on psychophysical thresholds. (**C**) Main axes: Markers represent M.A.P. estimates and transparent patches the corresponding 95% confidence intervals of threshold parameters fit to individual animals’ performance on control (y axis) versus mild cooling blocks (x axis). Single animals contribute one data point of each color. Top-right inset: Distribution of threshold differences between the mild cooling and control conditions (mean ± s.e.m.). (**D**) Left: Distributions of threshold dilation as a function of induced temperature changes. Markers linked by solid black lines represent individual animal threshold dilations. Right: Distribution of threshold stretch. Markers represent individual animals, and their size and color denote bootstrapped significance. (**E**) Same as (A), but in the *fixation* task variant. (**F-H**) Analogous to (B-D), but in the *fixation* task variant (mean ± propagated s.e.m. across trials instead of animals in (F)).

**Extended Data Figure 6.**
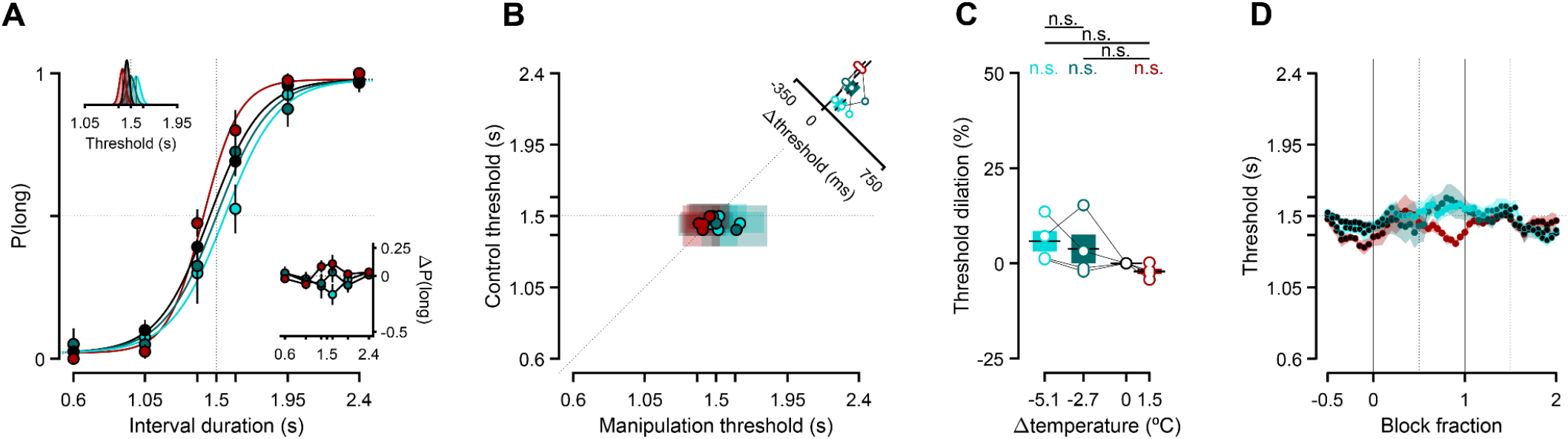
Manipulating M1 temperature did not produce discernible effects on timing judgments. (**A**) Average discrimination performance in the *fixation required* version of the interval discrimination task at the onset of M1 temperature manipulations. Main axes: psychometric functions fit to cross-animal averages (N = 4) of temperature-split psychophysical data, respectively shown as solid lines and markers of matching color (mean ± s.e.m.). Bottom-right inset: Average differences in proportion of long choices from each manipulation condition to control (mean ± propagated s.e.m.). Top-left inset: Marginal posterior distributions of the threshold parameter for each condition’s psychometric fit. Solid black lines represent the M.A.P. point estimates implicit in the fits shown in the main axes. (**B**) Animal-split discrimination behavior at the onset of M1 temperature manipulations. Main axes: Markers represent M.A.P. estimates and transparent patches the corresponding 95% confidence intervals of threshold parameters fit to individual animals’ performance on control (y axis) versus manipulation blocks (x axis). The identity line is plotted as a diagonal line. Inset: Distribution of threshold differences between manipulation and control conditions. Markers represent individual animal differences, bars and error bars are animal means and s.e.m. (**C**) Effect of motor cortical temperature manipulations on psychophysical threshold. Markers represent individual metric dilations, linked within animals by thin solid black lines. Boxplots show animal means (horizontal thick black lines) and s.e.m. (colored bars). (**D**) Threshold dynamics at the onset of M1 temperature manipulations, aligned to and across block transitions. Condition-split cross-animal average thresholds were computed using trials that fell into a sliding window lasting 90 s (half the block duration) that was swept from the preceding to the succeeding control blocks in increments of 9 s. Each marker corresponds to one sweep, and its color shading denotes the fraction of that sweep’s window that was inside a control block (with black markers corresponding to 100% control trials), and by extension its complement that was inside the manipulation block (with pure manipulation colors corresponding to 100% manipulation trials).

**Extended Data Figure 7.**
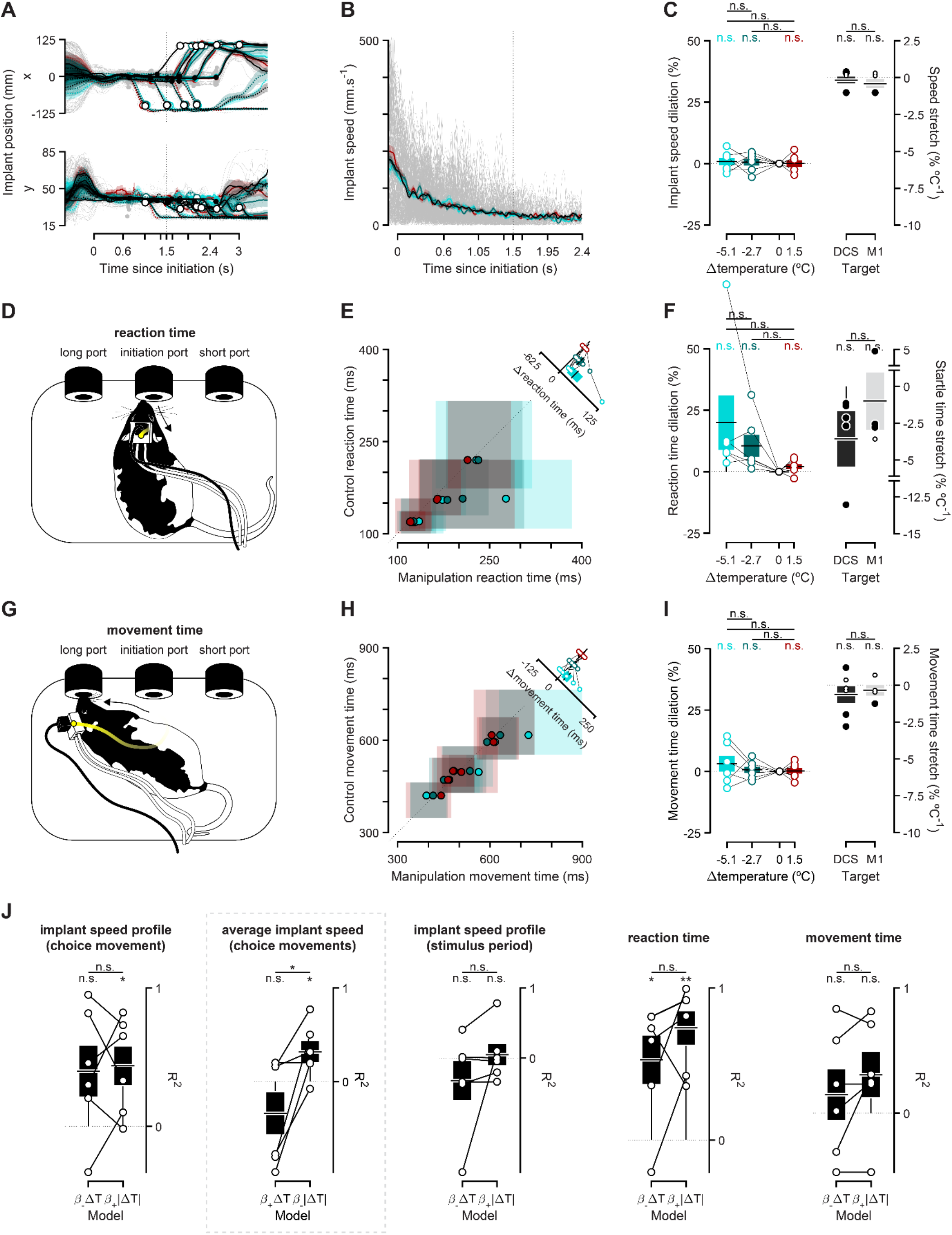
Temperature did not produce monotonic effects on movement during stimulus presentation, latency to initiate, or time to execute choice movements. (**A**) Horizontal (top) and vertical (bottom) coordinates of implant position aligned to stimulus onset (representative example animal). Dashed (solid) lines correspond to short (long) choices with individual control trials ghosted in the background, and condition-split averages on top. Filled and open markers show position at reaction and choice, respectively. (**B**) Same data as in (A), combined into an overall speed metric. (**C**) Implant speed dilation for DS animals (left) and stretch for DS and M1 animals (right). (**D**) Schematics highlighting the task epoch in between stimulus offset and the initiation of the choice movement (reaction time). **(E)** Main axes: Markers represent the median and transparent patches the corresponding i.q.r. of individual animals’ reaction times on control (y axis) versus manipulation blocks (x axis). Single animals contribute one data point of each color. The identity line is plotted as a diagonal line. Its thick solid portion highlights the region of the main axes that is shown in the bottom-right inset axes. Top-right inset: Distribution of reaction time differences between manipulation and control condition. Markers represent individual animal differences, bars and error bars are animal means and s.e.m. (**F**) Left: Distributions of median reaction time dilations as a function of induced temperature changes. Markers represent individual reaction time dilations, linked within animals by thin solid black lines. Right: Distribution of median reaction time stretches. Markers represent individual animals, and their size and color denote bootstrapped significance. (**G**-**I**) Same as (D-F), but for the task epoch in between the initiation of the choice movement and the moment that choice is registered (movement time). (**J**) Comparison between 2 one-parameter models of how temperature affected all movement-related metric dilations reported. On each panel, the goodness of fit of the monotonic (left) and the non-monotonic (right) model, as measured by its coefficient of determination (R^2^), is shown for individual animals (open markers) and cohorts (mean ± s.e.m.). Note that the *β*coefficients were constrained to positive or negative values, as indicated by their subscripts. The highlighted panel (dashed gray rectangle) is reproduced from Fig. 5f (right panel).

**Extended Data Figure 8.**
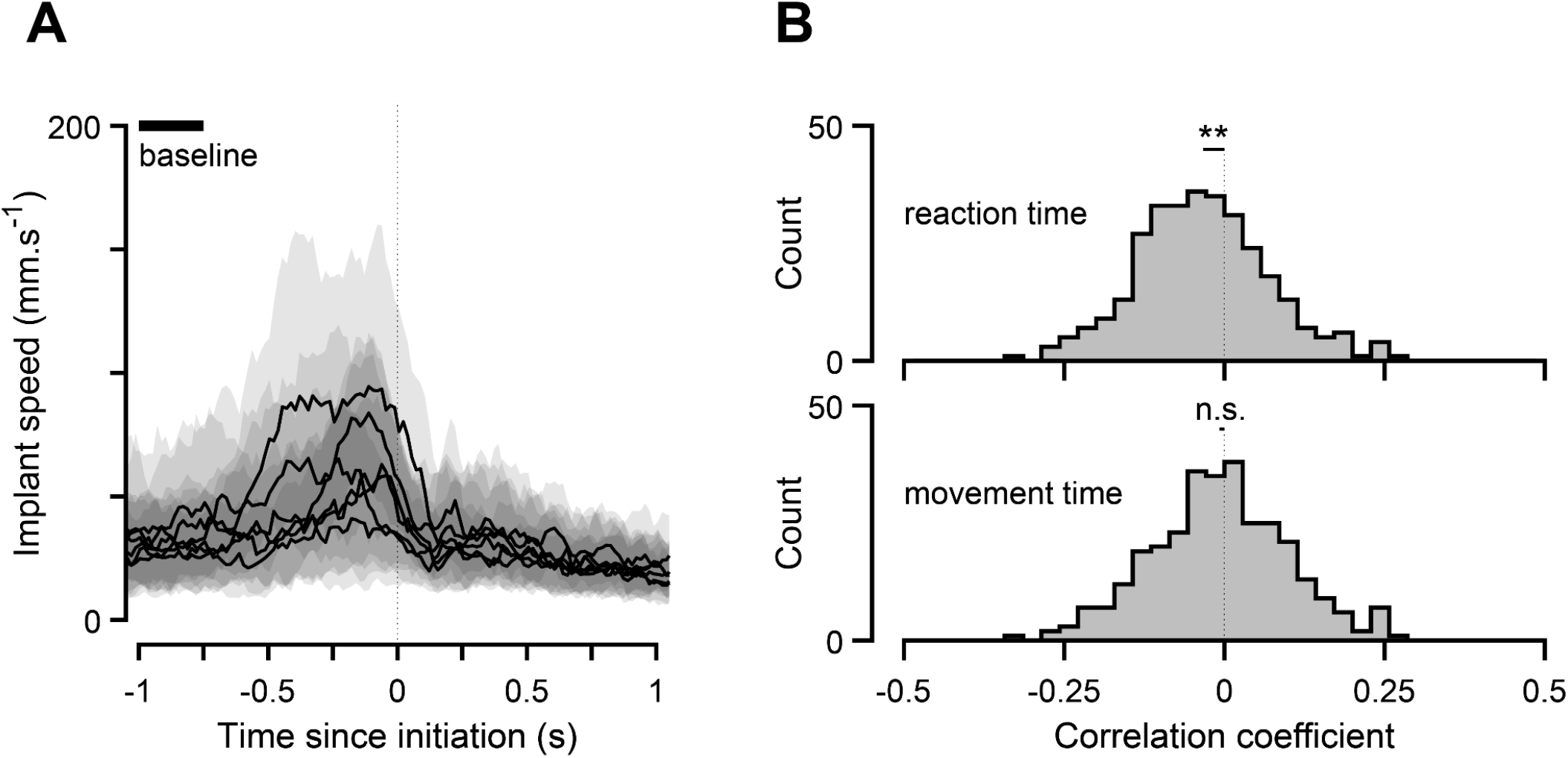
Baseline DS firing rate correlates with reaction but not movement times in the fixation version of the interval discrimination task. (**A**) Average implant speeds aligned to stimulus onset recorded during control blocks in the *fixation* version of the interval discrimination task (N = 6). Solid black lines and shaded gray patches represent individual animal medians and i.q.r., respectively. (**B**) Distribution of correlation coefficients between baseline firing rates of individual striatal neurons recorded during the *fixation* task variant (N = 483 neurons) and subsequent reaction (top) or movement times (bottom).

**Extended Data Figure 9.**
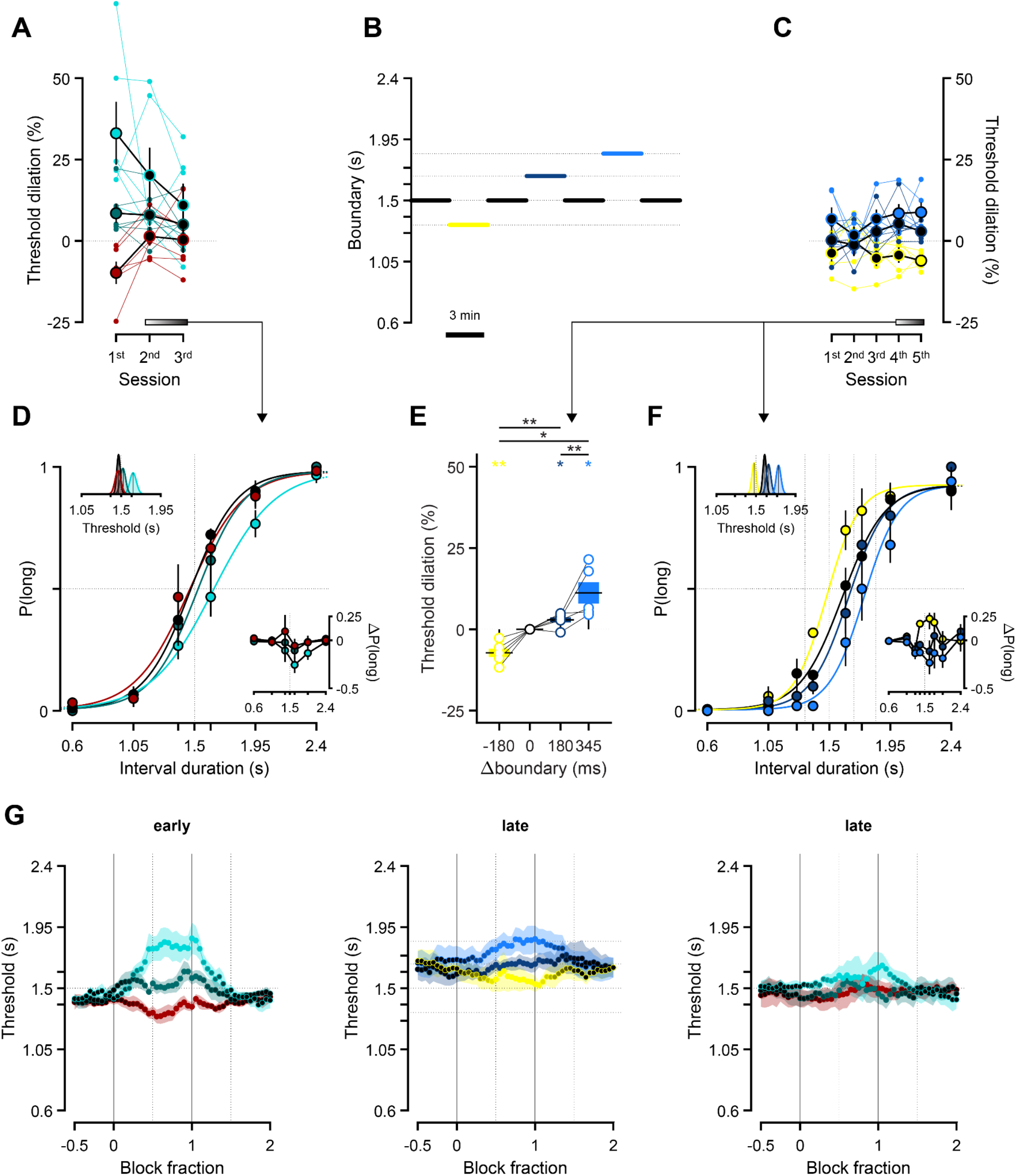
Animals adapted their behavior to both temperature and category boundary manipulations. (**A**) Threshold dilation across the first three temperature manipulation sessions for the striatal cohort shown in Fig. 2 (N = 6). Small markers and thin lines linking them refer to threshold dilations for individual animals. Larger markers correspond to cross-animal averages (mean ± s.e.m.), and their facecolor being any other than black indicates that the underlying dilation distribution was significantly shifted from zero (p<0.05, one-sample two-tailed t-test). The gradient bar and arrow symbolize the uneven contribution of the last two sessions to the data pool shown in (D), with the last session contributing the most. (**B**) Time course of the boundary manipulation experiment. The thin horizontal dotted lines represent the four categorical boundaries animals experienced in these sessions (i.e., boundary changes followed the same rules as the temperature manipulation experiments: a control-manipulation-control 3-min block design with boundaries drawn at random and without replacement from the set *B* = {1.32, 1.5, 1.68, 1.85} s until exhaustion, at which point the set was replenished and the sampling process resumed). The color scheme introduced in this panel is preserved throughout the figure. (**C**) Same as (A), but for the first five boundary manipulation sessions (N = 5 rats). (**D**) Average discrimination performance on the last and second to last sessions of striatal temperature manipulations. Main axes: psychometric functions fit to cross-animal averages of temperature-split psychophysical data, respectively shown as solid lines and markers of matching color (mean ± s.e.m.). Bottom-right inset: Average differences in proportion of long choices from each manipulation condition to control (mean ± propagated s.e.m.). Top-left inset: Marginal posterior distributions of the threshold parameter for each condition’s psychometric fit. Solid black lines represent the M.A.P. point estimates implicit in the fits shown in the main axes. (**E**) Distributions of percentage change in threshold relative to control (dilation) as a function of which categorical boundary was enforced. Markers represent individual threshold dilations, linked within animals by thin solid black lines. (**F**) Same as (D), but for the last two days of boundary manipulations, with all boundaries in our manipulation set as dotted vertical dashed lines. (**G**) Threshold dynamics aligned to and across block transitions early and late during DS temperature manipulations (left and right, respectively), and late during boundary manipulations (middle). Condition-split cross-animal average thresholds were computed using trials that fell into a sliding window lasting 90 s (half the block duration) that was swept from the preceding to the succeeding control blocks in increments of 9 s. Each marker corresponds to one sweep, and its color shading denotes the fraction of that sweep’s window that was inside a control block (with black markers corresponding to 100% control trials), and by extension its complement that was inside the manipulation block (with pure manipulation colors corresponding to 100% manipulation trials).

